# Beyond Respiration: Heme A Synthase (CtaA) Orchestrates Hydrogen Sulfide Production and Virulence via Metabolic Reprogramming in *Staphylococcus aureus*

**DOI:** 10.64898/2026.08.03.742422

**Authors:** Jiahui Li, Mu He, Wanwan Hou, Zengfeng Zhang, Zifeng Mai, Xiaorong Tian, Nan Zhong, Shimo Kang, Chunlei Shi

**Author notes:** Correspondence to Prof. Chunlei Shi.

## Abstract

Endogenous hydrogen sulfide (H_2_S) serves as a pivotal gasotransmitter conferring antimicrobial resistance and shielding bacteria from oxidative stress. While cystathionine γ-lyase (CSE)-dependent H_2_S production is critical for *Staphylococcus aureus* survival, the upstream regulatory nodes linking metabolic status to H_2_S-mediated redox defense remain elusive. Here, we identify heme A synthase (CtaA) as a master regulator that couples respiratory metabolism with H_2_S biosynthesis, specifically under glucose-depleted conditions mimicking host niches such as abscesses and phagosomes. Using a Himar1 transposon screen followed by a high efficiency detection method, we demonstrate that CtaA deficiency precipitates a catastrophic collapse in H_2_S levels (<10% of wild-type), leading to severe virulence and antimicrobial resistance changes. Specifically, the Δ*ctaA* mutant exhibits attenuated hemolytic activity and significantly reduced virulence in a *Galleria mellonella* model, despite displaying paradoxical resistance to specific antimicrobials.

Mechanistically, CtaA deletion triggers a maladaptive metabolic reprogramming characterized by the downregulation of the L-cysteine transporter TcyP and dysregulation of arginine metabolism, which collectively impair the bacterium’s capacity to produce endogenous H_2_S to defense oxidative stress. Notably, this redox vulnerability is linked to altered endogenous nitric oxide (NO) dynamics, suggesting a disrupted H_2_S-NO crosstalk essential for stress adaptation. Our findings elucidate a novel metabolic-redox axis where CtaA governs H_2_S homeostasis to counteract host-imposed oxidative stress. Targeting the CtaA-H_2_S axis represents a promising therapeutic strategy to sensitize *S. aureus* to host immune clearance by dismantling its critical redox shield.

## 1. Introduction

Hydrogen sulfide (H_2_S) has transcended its historical reputation as a toxic byproduct to emerge as a ubiquitous gasotransmitter that governs cellular redox homeostasis across all domains of life [1–3]. In bacterial pathogens, endogenous H_2_S serves as a critical defense shield[4], scavenging reactive oxygen (ROS) generated by host immune cells and conferring tolerance to antimicrobial stress[5, 6]. While the enzymatic machinery for H_2_S biosynthesis—primarily cystathionine γ-lyase (CSE), cystathionine β-synthase (CBS) and 3-mercaptopyruvate sulfurtransferase (3-MST)—is well conserved [1, 6], the upstream regulatory networks that dynamically tune H_2_S production in response to metabolic cues remain largely elusive. This gap in knowledge is particularly pronounced in *Staphylococcus aureus*, a formidable pathogen that must precisely calibrate its redox defenses to survive within hostile, nutrient-depleted host niches such as abscesses and phagolysosomes [7, 8].

Current understanding of bacterial H_2_S regulation is predominantly framed within the context of sulfur metabolism. Canonical regulators like CymR in *S. aureus* [9] and CysB in *Escherichia coli* [10] modulate cysteine uptake and assimilation, thereby indirectly influencing substrate availability for H_2_S synthesis [11]. Furthermore, links between iron homeostasis (via Fur) and H_2_S-mediated oxidative stress tolerance have been established, highlighting the integration of gas signaling with metal metabolism [10, 12, 13]. However, these “sulfur-centric” or “metal-centric” models fail to account for how global bioenergetic states—specifically the functional status of the respiratory chain—retrograde-regulate gasotransmitter production. Given that H_2_S can reversibly inhibit cytochrome *c* oxidase and modulate respiratory flux in mitochondria and bacteria [14–16], a reciprocal regulatory loop likely exists: does the integrity of the respiratory chain itself dictate the capacity for H_2_S-mediated redox defense?

The aerobic respiratory chain of *S. aureus* presents a compelling, yet unexplored, node for such regulation. The bacterium relies on two terminal oxidases: the high-affinity cytochrome *bd* (CydAB) and the high-efficiency cytochrome *aa_3_* (QoxABCD) [17]. The biogenesis of the latter is strictly dependent on heme A, a specialized cofactor synthesized from heme O by the enzyme heme A synthase (CtaA) [18]. CtaA is traditionally viewed as a mere assembly factor essential for aerobic respiration and ATP generation under oxygen-rich conditions[19]. Yet, emerging paradigms in redox biology suggest that respiratory complexes act as signaling hubs that sense metabolic stress and orchestrate broader adaptive responses [20]. We hypothesized that CtaA, by governing the maturation of the cytochrome *aa_3_* complex, might serve as a critical metabolic switch that links respiratory efficiency to the production of endogenous H_2_S production in *S. aureus*.

This hypothesis addresses a critical translational challenge. Direct inhibition of CSE has been proposed as a strategy to sensitize *S. aureus* to antibiotics [6]; however, the high structural conservation of PLP-dependent enzymes between hosts and bacteria raises significant concerns about off-target toxicity and metabolic acidosis [21, 22]. Identifying upstream, bacteria-specific regulatory nodes that control H_2_S homeostasis—such as the interface between heme biosynthesis and gas signaling—offers a more promising therapeutic avenue. Such targets could dismantle the bacterial redox shield without compromising host physiology. Despite the established role of CtaA in respiratory competence, its potential involvement in regulating non-respiratory functions, specifically the homeostasis of gasotransmitters, has never been investigated.

In this study, we uncover a previously unrecognized function of CtaA as a master regulator of endogenous H_2_S production in *S. aureus*. Using a Himar1 transposon screen combined with transcriptomic profiling and biochemical assays, we demonstrate that CtaA deficiency precipitates a catastrophic collapse in H_2_S levels, driven by a profound metabolic reprogramming of cysteine and arginine pathways. We reveal that under glucose starvation—a condition mimicking the host environment—CtaA acts as a central nexus linking heme A biosynthesis, nitric oxide (NO) dynamics, and H_2_S-mediated antioxidant defense. Mechanistically, we show that CtaA loss disrupts the H_2_S-NO crosstalk and impairs the expression of key sulfur transporters, leading to heightened susceptibility to oxidative stress despite enhanced resistance to certain antimicrobials. These findings redefine CtaA from a simple respiratory assembly factor to a pivotal conductor of the metabolic-gasotransmitter axis, providing new mechanistic insights into how *S. aureus* orchestrates its virulence and stress resilience. This work highlights the CtaA-H_2_S axis as a potent, bacteria-specific target for next-generation antimicrobial therapies aimed at disrupting bacterial redox homeostasis.

## 2. Materials and methods

### 2.1 Bacterial strains, plasmids, and media

The bacterial strains and plasmids used in this study are listed in Table 1. Mutant D12 is an endogenous H_2_S production-deficient mutant isolated from a *Himar1* transposon insertion mutant library of strain ATCC BAA1717, established as the workflow shown in Fig. 1. *Escherichia coli* JTU006 was used for plasmid methylation modification. For *E. coli*, bacterial cultures were incubated in lysogeny broth (LB) medium with shaking at 200 rpm or on LB agar plates at 37°C. For *S. aureus*, strains were grown in tryptic soy broth (TSB) with shaking at 200 rpm or passaged on tryptic soy agar (TSA) at 37°C. When necessary, the culture media were supplemented with antimicrobials (100 µg/mL ampicillin for *E. coli* and 10 µg/mL chloramphenicol for *S. aureus*).

**Fig. 1.**
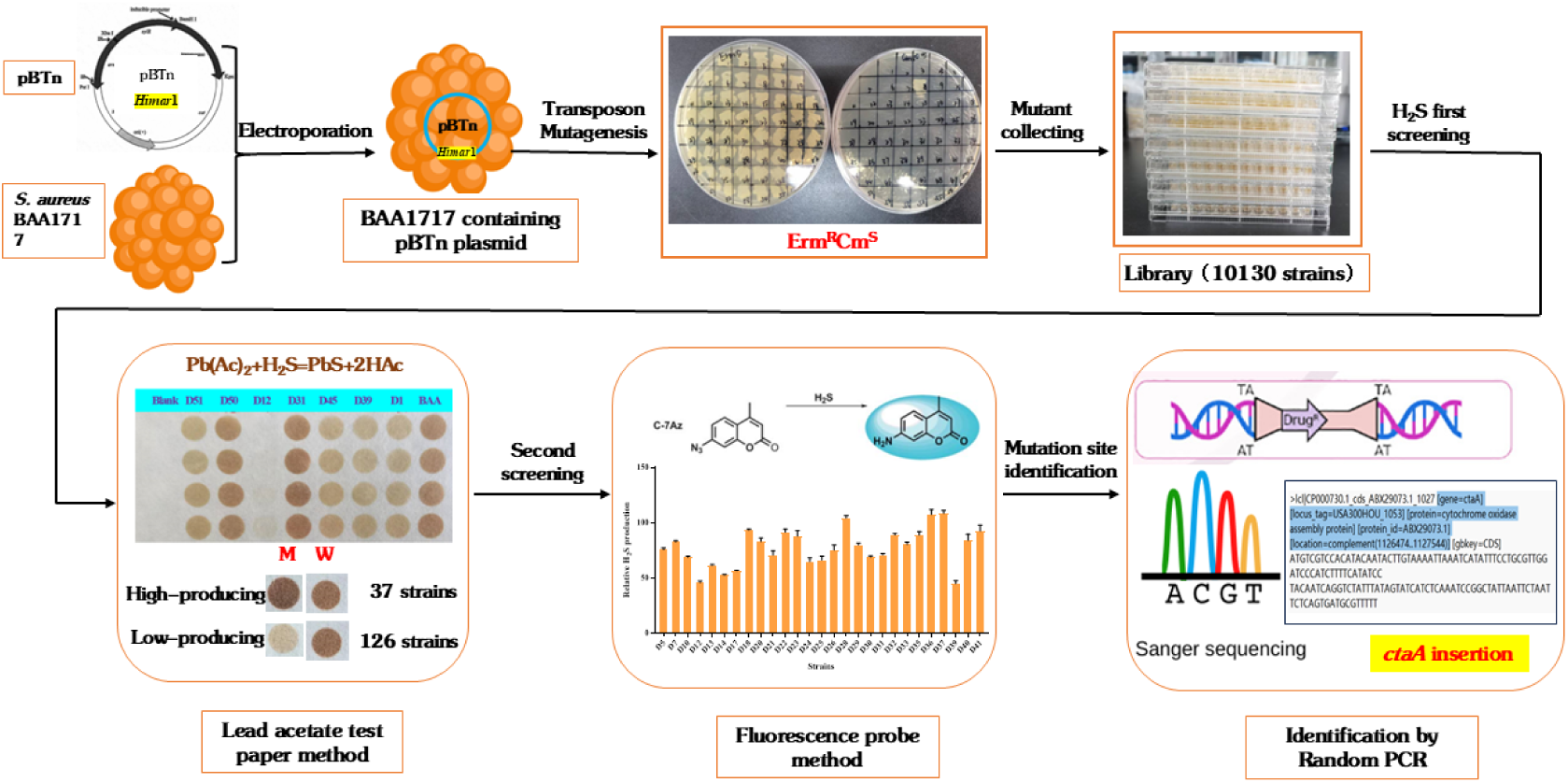
Flow chart for screening H_2_S low-producing mutant D12 by transposon insertion mutation.

**Table 1.**
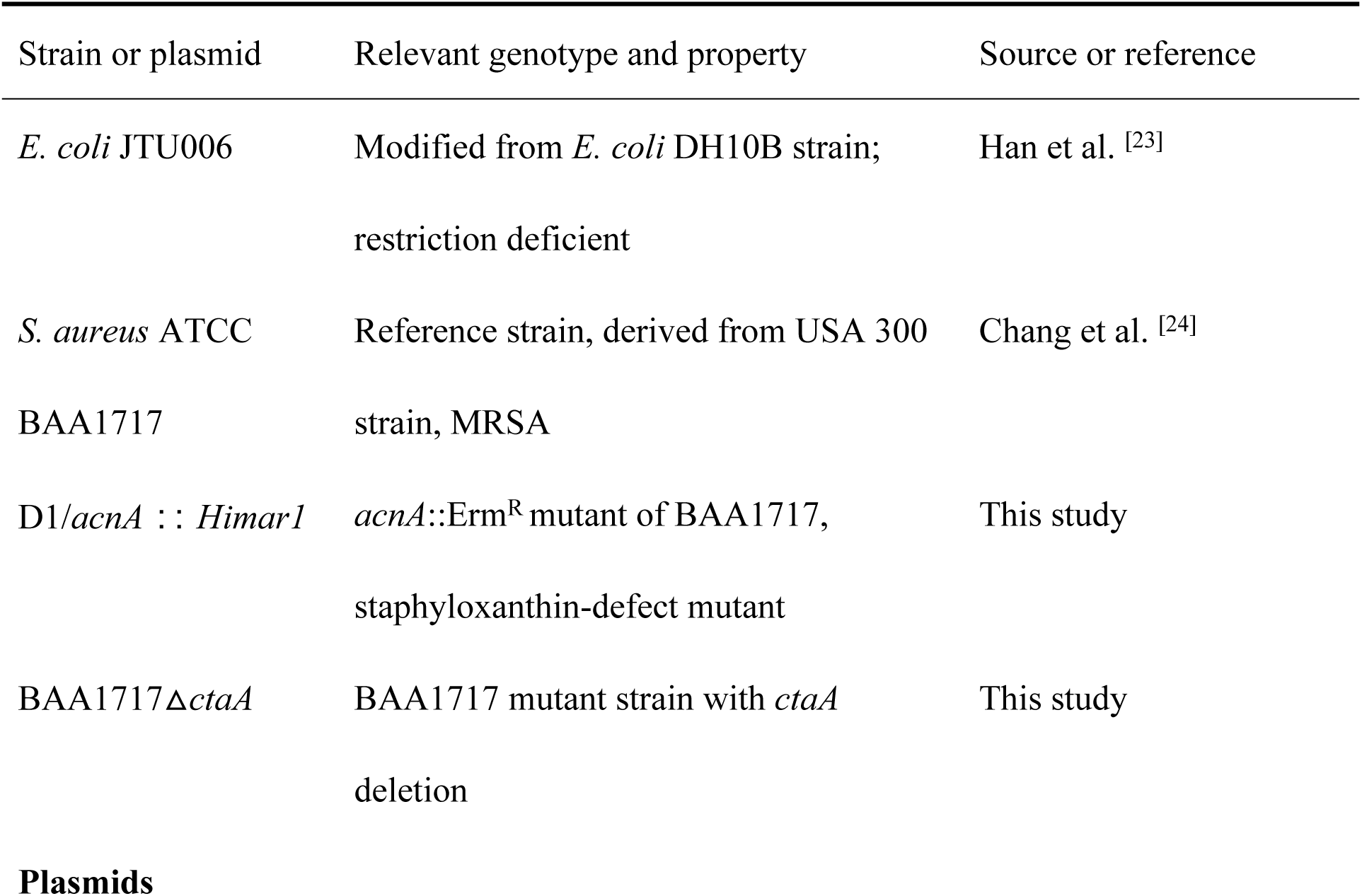
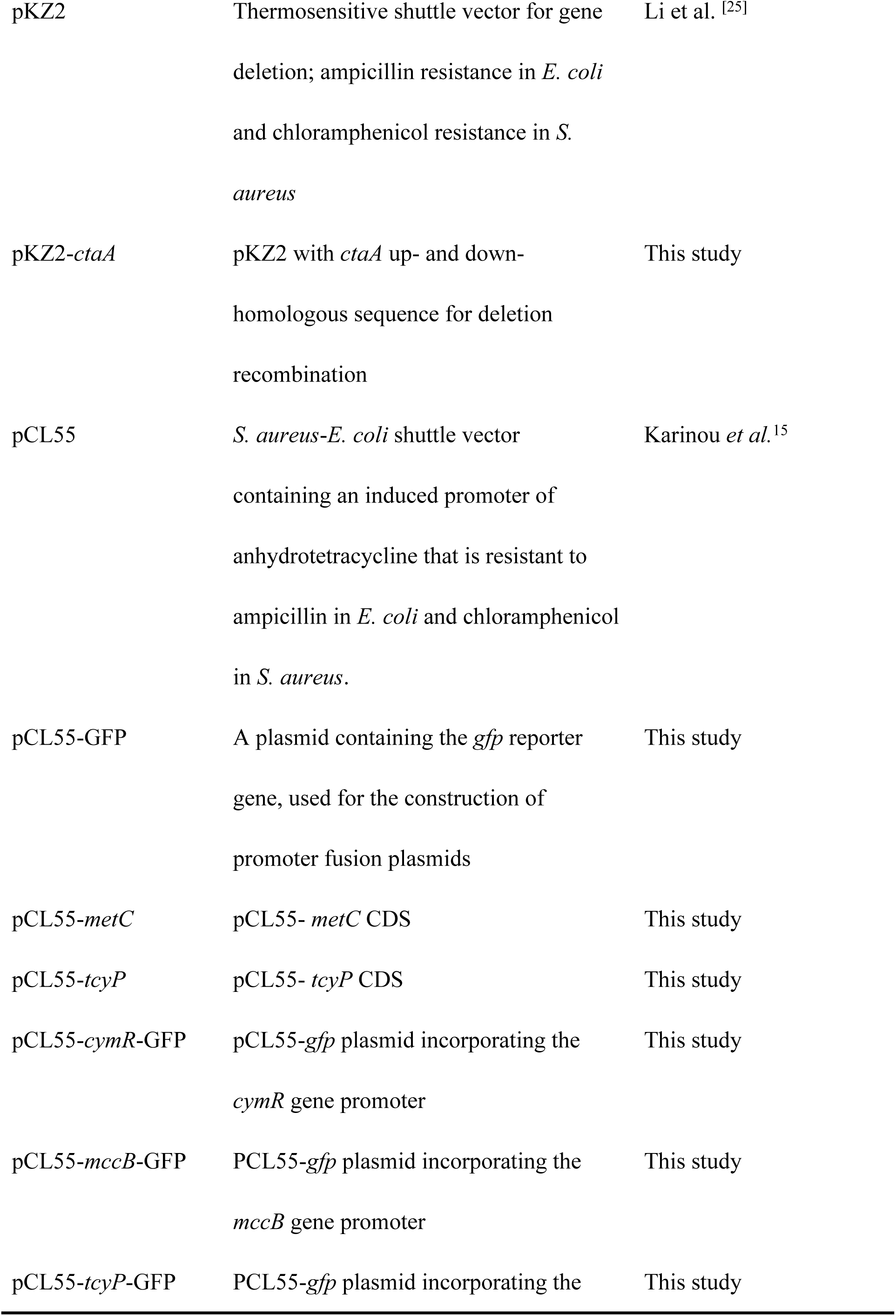
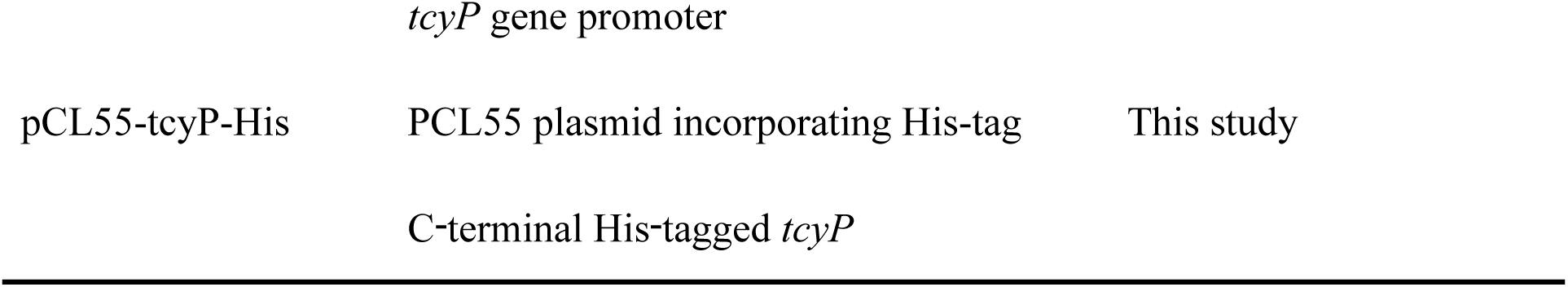
Strains and plasmids used in this study.

### 2.2 Gene deletion

The *ctaA* gene of strain BAA1717 was deleted by the method described previously [26]. The upstream and downstream DNA fragments of the *ctaA* gene were amplified by PCR. The homology arm fragment was then generated through overlap extension PCR. The resulting PCR products were cloned into the pKZ2 vector. The resultant plasmids were subsequently transformed into *E. coli* JTU006 for plasmid methylation modification. The modified plasmids were electroporated into strain BAA1717 to achieve *ctaA* gene knockout. Primers used for gene deletion are listed in Supplementary Table S1.

### 2.3 Construction of complementation and overexpression strains

For complementation and overexpression analyses, the coding sequences of the *metC* and *tcyP* genes, including their native start codon regions, were amplified from genomic DNA of the MRSA strain BAA1717. Each fragment was subsequently cloned into the plasmid vector pCL55, generating the recombinant plasmids pCL55-*metC* and pCL55-*tcyP*. These plasmids were then separately transformed into both the wild-type (WT) strain and the △*ctaA* mutant strain for complementation and overexpression assays, following the procedures outlined in the previous section. All primers used for plasmid construction are listed in Table S1.

### 2.4 Growth Curve assays

The growth curve was determined to evaluated the effect of *ctaA* gene deletion on the growth of *S. aureus*. In brief, an overnight bacterial culture was diluted to approximately 5×10^6^ CFU/mL. Aliquots (250 μL) of these suspensions were added to the wells of a 100-well honeycomb plate, with sterile TSB solution serving as a background control. The plate was incubated at 37℃ for 24 h, with OD_600_ measurements taken hourly to generate the bacterial growth curves.

### 2.5 H_2_S detection

The lead acetate detection method was modified to measure H_2_S production from different strains [6]. A 2% Pb-acetate-saturated paper piece was moistened and affixed above a 96-well plate containing bacterial suspensions with 200 µM L-cysteine. The plate was then sealed and incubated at 37℃ for 18 h with shaking at 200 rpm. The stained paper pieces were subsequently scanned and quantified with the ImageJ software (v1.53).

To investigate how *ctaA* affects endogenous H_2_S production, we combined lead acetate paper assays based on transcriptomics with verification by exogenous addition of key substrates or inhibitors. After culturing the strains to the logarithmic phase, the OD_600_ of the bacterial cultures was normalized. Various chemical compounds were then added to the culture medium and incubated. Endogenous H_2_S production under each treatment condition was quantified using the lead acetate paper method, as described above. The final concentrations of additives were as follows: glucose, 0–0.5%; zinc protoporphyrin (ZPP), 0–200 μM; FeCl_3_, 0–200 μM; arginine, 0–100 mM; NaNO₂, 0–8 mM; and NG-Nitro-L-arginine Methyl Ester (L-NAME), 0–8 mM.

For the measurement of dissolved H_2_S, a fluorescent-based detection method was employed [27]. In brief, overnight bacterial cultures were washed twice with phosphate-buffered saline (PBS) and adjusted to an OD_600_ of 1.0. The washed cells were then incubated for 2 h in PBS containing 100 µM of the 7-azido-4-methylcoumarin (AzMC) fluorescent probe (Sigma-Aldrich, CAS 95633-27-5, Saint Louis, USA), which selectively reacts with H_2_S to form a fluorescent compound. Fluorescence measurements were taken using a microplate reader (Molecular Devices, Sunnyvale, USA) with an excitation wavelength of 365 nm and an emission wavelength of 450 nm.

### 2.6 Staphyloxanthin production detection

The methanol extraction method was used to quantify staphyloxanthin production in *S. aureus* [24]. In brief, WT and △*ctaA* strains were cultured in TSB at 37°C for 48 h. The cells were then harvested by centrifugation at 8,000 rpm for 5 min at 4°C and washed three times with PBS. Staphyloxanthin was extracted by adding methanol, followed by incubation at 55°C for 3 h. After incubation, the samples were centrifuged at 6,000 rpm for 2 min at 4°C, and the supernatant was collected. The staphyloxanthin content was determined by measuring the optical density at 462 nm using a Sunrise^®^ spectrophotometer (Tecan, Mannedorf, Switzerland).

### 2.7 Hemolysis assays

Hemolysis activity was evaluated using sheep red blood cells, as described by Zheng *et al*. [28]. Briefly, *S. aureus* strains were incubated at 37℃ for 18 h with shaking at 200 rpm. The cultures were then centrifuged at12000rpm for 10 min at 4℃, and the supernatants containing hemolysin were collected. Aliquot (250 μL) of the supernatants were added to a solution containing 1% fresh sheep red blood cells. The degree of hemolysis was quantified by measuring the absorbance at 540 nm. Complete lysis of red blood cells, achieved using 1% Triton X-100, served as the positive control (100% hemolysis), while TSB served as the negative control (0% hemolysis).

### 2.8 Biofilm formation assay

Biofilm quantification was performed using the crystal violet assay [29]. Briefly, an overnight culture of *S. aureus* was harvested and adjusted to OD_600_ of 1.0 using TSB medium. The bacterial suspension was then transferred to 96-well plates and incubated statically for 24 hours at 37°C. After incubation, planktonic wells were removed by washing with PBS. The adherent biofilm was stained with 0.1% crystal violet for 20 min, after which excess dye was removed by thorough washing. The bound crystal violet was subsequently dissolved in ethanol, and the biofilm formation was quantified by measuring the absorbance at 595 nm.

### 2.9 Evaluation of infectivity using a *Galleria mellonella* model

The virulence of the WT and △*ctaA* mutant strains was evaluated using an *in vivo* infection model in *Galleria mellonella* larvae, as previously described [30]. Larvae weighing approximately 200 mg were randomly assigned to experimental groups (n = 10 per group). Each larva was infected by injecting 10 μL of bacterial suspension containing either the WT or △*ctaA* mutant strain at a concentration of 1.0 × 10⁵ CFU/ml in PBS into the last left posterior leg. After injection, the larvae were incubated at 37°C and monitored for survival over a 5-day period post-infection, with mortality recorded daily. This protocol enabled a comparative analysis of the survival outcomes and replication capacity of the △*ctaA* mutant and WT strains within a live host environment.

### 2.10 Antimicrobial susceptibility test

Antimicrobial susceptibility testing of *S. aureus* was conducted using the microdilution method as outlined in the CLSI guidelines (2018). Briefly, bacterial cultures were adjusted to a concentration of 5×10^6^ CFU/mL and added to wells containing serial dilutions of various antimicrobials in TSB. After incubation at 37°C for 24 h, the minimum inhibitory concentration (MIC) was determined by identifying the lowest concentration of the antimicrobial that inhibited visible bacterial growth.

### 2.11 SEM assay

To evaluate the morphology of the strains, field-emission scanning electron microscopy (FESEM) was utilized [31]. Bacterial cells were fixed with 2.5% (v/v) glutaraldehyde overnight at 4°C, dehydrated through a graded ethanol series, and subsequently gold-coated to enhance conductivity for electron microscopy. FESEM imaging was performed using an SEM (TM3030, Hitachi, Tokyo, Japan) at an accelerating voltage of 15 kV, enabling high-resolution observation of cell surface features, including cellular arrangement and surface topography. Maximum cell diameter was measured as the average of 60 randomly selected cells by ImageJ (v1.53).

### 2.12 TEM assay

The morphology and ultrastructure of *S. aureus* strains were analyzed using a transmission electron microscope (TEM) [32]. Bacterial samples were initially fixed in 2.5% glutaraldehyde for 2 h at 4°C, followed by post-fixation in 1% osmium tetroxide for 1 h. Dehydration was performed using a graded ethanol series (30%, 50%, 70%, 90%, and 100%), after which the samples were embedded in epoxy resin and polymerized at 60°C. Thin sections (70 nm) were prepared using an ultramicrotome and stained with uranyl acetate and lead citrate. High-resolution images of the detailed structural features, including the cell wall structure of the *S. aureus* strains, were captured using TEM at an accelerating voltage of 120 kV. Cell wall thickness was measured as the average of 20 randomly selected cells by ImageJ (v1.53).

### 2.13 RNA-Seq analysis

Overnight cultures of the WT and △*ctaA* mutant strains, incubated with 0.25% glucose or without glucose in TSB, were adjusted to an OD_600_ of 1.0. For RNA extraction, the cultures were harvested by centrifugation at 4°C. Total RNA was extracted by an Invitrogen Trizol extraction kit (15596-026, California, USA), following the method described by Hu *et al*. [33]. Ribosomal RNA was removed using the Zymo-Seq RiboFree Total RNA Library Kit (R3003, California, USA). The sequencing library was then sequenced on the NovaSeq 6000 platform (Illumina, San Diego, USA) by Shanghai Personal Biotechnology Co. Ltd. Differential expressed genes (DEGs) were identified using the edgeR software package with a log_2_ fold change (>1.0) and a *P*-value (< 0.05). The RNA-seq experiment was performed with three independent biological replicates.

### 2.14 RT-qPCR qualification

Total RNA of *S. aureus* cells were extracted using kit (R701-01, Vazyme, Nanjing, China) as previously described [34]. The extracted RNA was reverse-transcribed into cDNA using a PrimeScript RT reagent kit with gDNA eraser (R323-01, Vazyme, Nanjing, China). Subsequently, RT-qPCR was performed using SYBR Green PCR Master Mix (Q711-03, Vazyme, Nanjing, China) on a CFX96 Real-Time System (Eppendorf, Hamburg, Germany). Gene expression changes between strains were assessed using the ΔΔCt method, with the *gyrB* gene serving as the housekeeping gene. Fold changes were calculated using the 2^-ΔΔCt^ method.

### 2.15 L-cysteine content determination

The L-cysteine assay kit (BC0185, Solarbio, Beijing, China) was used to determine the intracellular L-cysteine concentration. L-cysteine reduces phosphotungstic acid to tungsten blue, which has an absorption peak at 600 nm. The L-cysteine content was determined by measuring the absorbance at this wavelength. In brief, 4 mL of overnight cultures were centrifuged at 8,000 rpm for 3 min. The cell pellets were then resuspended in 1 mL of extraction solution, mixed thoroughly, and sonicated in an ice bath (200 w, 3 s of ultrasound, 7 s of interval, total time of 5 min). After centrifugation at 11,000 rpm for 10 min at 4°C, the supernatant was collected and kept on ice for further analysis. The L-cysteine concentration was quantified according to the manufacturer’s instructions.

### 2.16 Construction of the GFP reporter plasmid

To construct the GFP reporter plasmid pCL55-target gene-GFP, the promoter region of each target gene (*cymR*, *mccB*, or *tcyP*), along with the first 18 bp of its coding sequence, was PCR-amplified from genomic DNA of *S. aureus* strain BAA1717 using gene-specific primers (Ptarget gene-F/Ptarget gene-R). The resulting PCR fragment was then ligated into the *Eco*RV-digested pCL55 vector via seamless cloning, yielding the corresponding reporter plasmid. Following sequence verification, each recombinant plasmid was first transformed into *E. coli* JTU006 for propagation and modification, and subsequently introduced into both the WT and Δ*ctaA* mutant strains of *S. aureus* by electroporation.

### 2.17 GFP reporter assays

*S. aureus* strains harboring the GFP reporter plasmid were grown overnight in the presence of 10 μg/mL chloramphenicol. The overnight cultures were harvested, washed twice with PBS buffer, and adjusted to an optical density (OD_600_) of 1.0 ± 0.05. For GFP fluorescence quantification, 200 μL of each sample was loaded in triplicate into a black 96-well microplate. Fluorescence was measured using a microplate reader (Molecular Devices, Sunnyvale, USA) with excitation at 485 nm and emission at 520 nm. To compare promoter activity between the WT and the Δ*ctaA* mutant strains, GFP fluorescence readings were first normalized to the cell density (OD_600_) of each sample, and then to the background fluorescence of the corresponding strain harboring the empty vector pCL55-GFP.

### 2.18 Western blot assays

To compare TcyP expression levels between the WT strain and the Δ*ctaA* mutant, a recombinant plasmid, pCL55-tcyP-His, was constructed. This plasmid encodes a C-terminal His-tagged TcyP fusion protein (predicted molecular weight ≈ 50 kDa) and was introduced into both the WT and Δ*ctaA* backgrounds. Whole-cell protein extracts were separated on 12% SDS-PAGE and transferred to a membrane for immunoblotting. His-tagged TcyP was detected using a Horseradish Peroxidase (HRP)-conjugated anti-His mouse monoclonal antibody (AF2879, Beyotime, Shanghai, China). The WT strain harboring the empty vector (lacking the His-tag, pCL55) was used as a negative control, while endogenous GroEL served as a loading control.

### 2.19 NO content determination

The NO content was determined using a NO detection assay kit (D799310, Sangon, Shanghai, China). Overnight cultures of the WT and △*ctaA* mutant strains were harvested by centrifugation at 8,000 rpm for 5 min at 4℃. The resulting cell pellets were washed twice with sterile PBS buffer and resuspended in extraction buffer. Cells were lysed and centrifuged at 12,000 rpm for 15 min at 4℃. The supernatant was collected and kept on ice for subsequent analysis. NO content was quantified according to the manufacturer’s protocol. Total protein concentration in the supernatant was measured using a Bicinchoninic acid (BCA) protein assay kit (C503061, Sangon, Shanghai, China), and aconitase activities were normalized to the total protein content of each sample.

### 2.20 Statistical analysis

Statistical significance was assessed using GraphPad Prism 8.0. Significant differences were defined through two-way ANOVA analysis and Student’s *t*-test with *\*P* < 0.05, and \*\**P* < 0.01 indicating significant and highly significant differences, respectively.

## 3. Results

### 3.1 Tn-Seq Screening for Genes Regulating Endogenous H_2_S Production in *S. aureus*

A random *Himar1* transposon insertion library was constructed in *S. aureus* ATCC BAA1717 to identify genes involved in endogenous H_2_S production. The library was generated by conjugating the donor strain *E. coli* JTU006, which harbors the *Himar1* transposon, with the recipient *S. aureus* ATCC BAA1717, resulting in approximately 10,000 mutants (Fig. 1). To screen for mutants with altered H_2_S production, we developed a high-throughput method using lead acetate test paper, which detects H_2_S through the formation of a black lead sulfide precipitate. The intensity of the black coloration was positively correlated with H_2_S concentration, with darker spots indicating higher levels of H_2_S production. By comparing the spot intensities between individual mutants and the WT strain, we successfully isolated mutants exhibiting either increased or decreased H_2_S production based on clearly distinguishable phenotypic differences (Fig.1).

Furthermore, the transposon insertion sites of 30 low-H_2_S-producing mutants and 12 high-H_2_S-producing mutants were determined by inverse PCR to identify the genes involved in H_2_S production. To elucidate the molecular mechanisms underlying the altered endogenous H_2_S production in mutants, random PCR mutagenesis was employed to pinpoint the disrupted genes. In the 30 low-H_2_S-producing mutants, the transposon insertions were detected in 16 distinct genes or intergenic regions (Table S2), including *acnA, ctaA, oppA1, gpsA*, and others. Among the 12 high-H_2_S-producing mutants, insertions were found in 9 different genes or intergenic regions (Table S3), such as *USA300HOU_2364*, *pbuX*, and *USA300HOU_2169*.

### 3.2 Identification of a regulator, *ctaA*, that positively controls H_2_S production, particularly under glucose-free conditions

Mutant strain D12 was identified by the method described above, as it exhibited an apparent reduction in H_2_S production compared to the WT strain using the lead acetate test paper method under no-glucose conditions (Fig. 2A). Random PCR analysis revealed that the *Himar1* transposon was inserted between positions 228 and 229 within the coding region of the *ctaA* gene, which encodes a cytochrome oxidase assembly protein known as heme A synthase (Fig. 1). In *S. aureus*, heme A synthase catalyzes the conversion of heme O to heme A, a critical step in the function of cytochrome a-type respiratory oxidases. To confirm the precise impact on H_2_S production, the fluorescence probe method was employed, demonstrating a reduction of over 90% in H_2_S generation in strain D12 under glucose-free conditions (Fig. 2B).

**Fig. 2.**
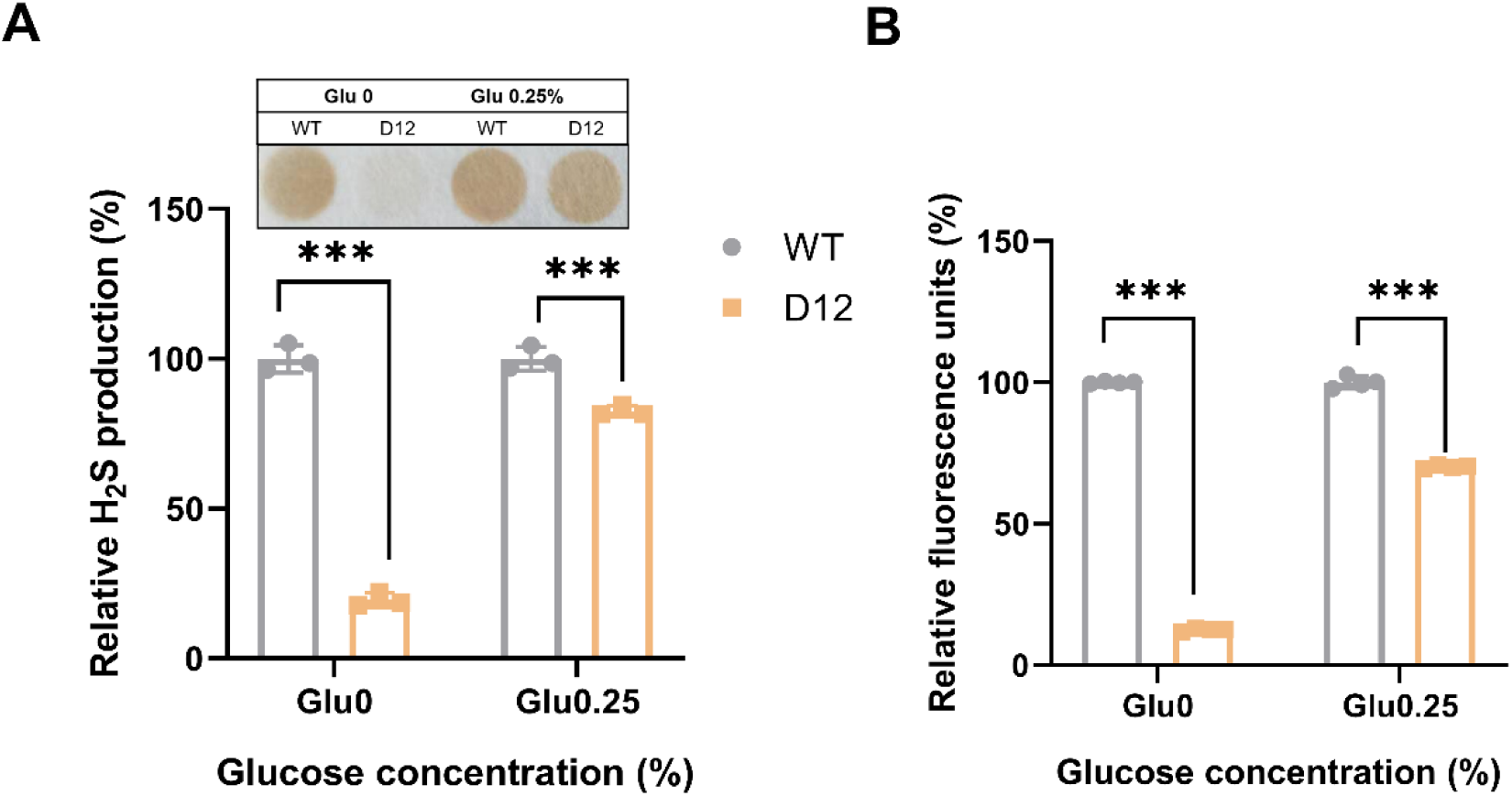
Identification of the H_2_S production regulator CtaA. (A) Lead acetate method to evaluate the ability of WT and D12 strains to produce H_2_S. (B) AzMC fluorescent probe method to determine the H_2_S production of WT and D12 strains. Data represents mean±SD from at least n = 3 biological replicates. \*\**P* < 0.01

### 3.2 Heme A synthase *ctaA* positively Regulates H_2_S production in *S. aureus* in a glucose-dependent manner

To further investigate the function of the *ctaA* gene, a deletion mutant strain, Δ*ctaA*, was generated by complete removal of the *ctaA* gene, and its capacity to produce endogenous H_2_S was assessed. Results demonstrated that H_2_S production was significantly reduced in the △*ctaA* mutant compared to the WT strain BAA1717 under glucose-free conditions, and was comparable to that of the *ctaA*-disrupted mutant D12 (Fig. 3A&3B). In the low-glucose range (0-0.25%), H_2_S production in the △*ctaA* mutant increased with rising glucose concentration, whereas H_2_S levels in the WT strain remained relatively stable. However, further glucose supplementation beyond 0.25% markedly suppressed H_2_S production in both strains, with complete inhibition observed at 0.5% glucose. These findings collectively indicate that CtaA functions as a glucose-dependent regulator of H_2_S production in *S. aureus*.

**Fig. 3.**
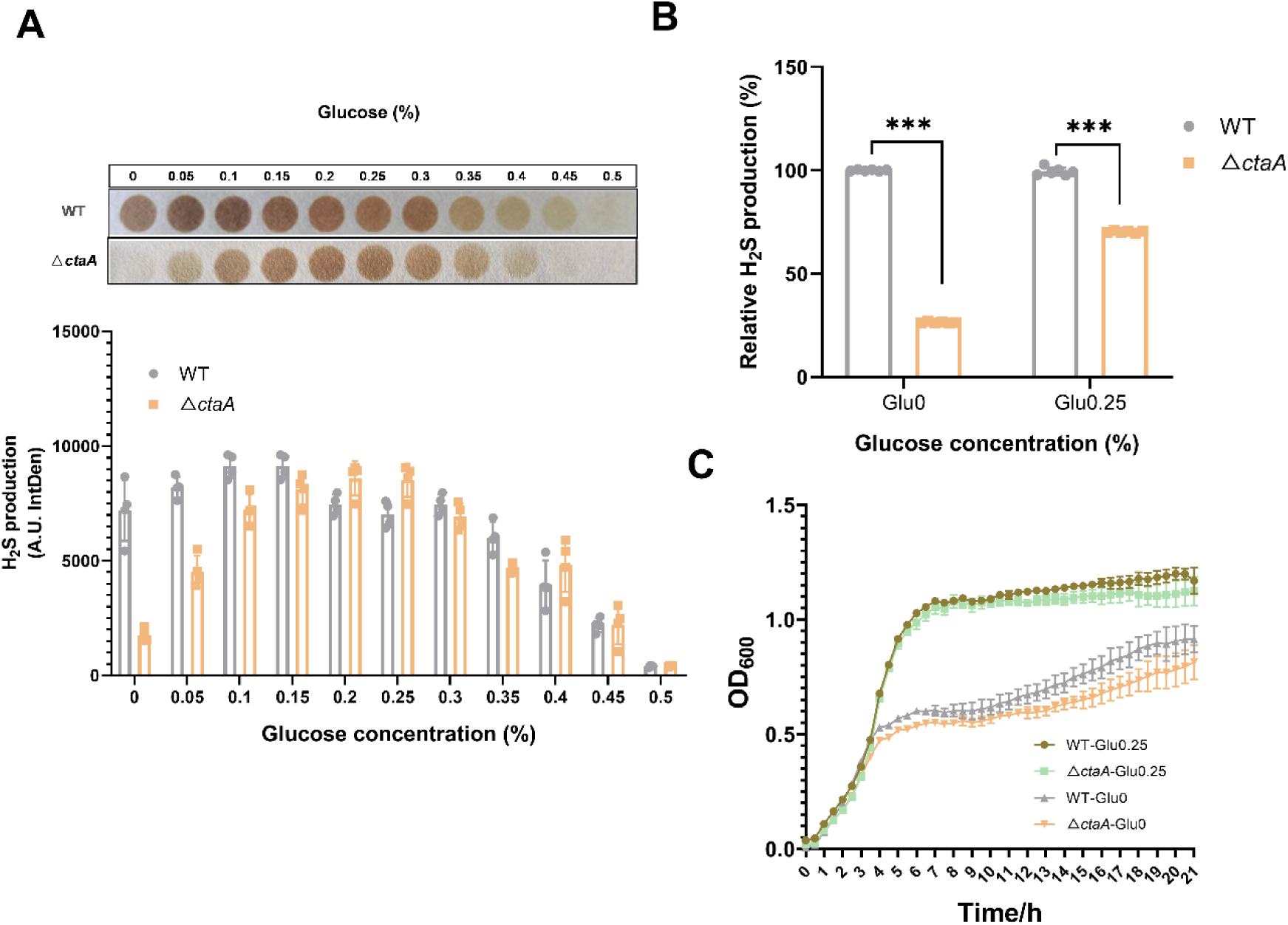
H_2_S production and growth curve determination of △*ctaA* mutant. (A) H_2_S production determination by Lead acetate method under different concentrations of glucose (0∼0.5%). (B) H_2_S production determination by AzMC fluorescent probe method under 0.25% glucose and glucose-free conditions. (C) Growth curve determination of the WT and △*ctaA* mutant strains under 0.25% glucose and glucose-free conditions. Data represents mean±SD from at least n = 3 biological replicates. \*\**P* < 0.01.

To determine whether reduced H_2_S production in the △*ctaA* mutant was attributed to impaired cell growth, growth curves of the WT and △*ctaA* strains were compared under both glucose-containing and glucose-free conditions. As shown in Fig. 3C, the growth rate of *S. aureus* was mildly inhibited in both conditions, with greater inhibition observed in the absence of glucose, suggesting that the △*ctaA* mutant is more sensitive to glucose deprivation than the WT strain. Given that the △*ctaA* mutant exhibited a substantial reduction in H_2_S production under glucose-free conditions, a minor decrease in growth rate is unlikely to be the primary cause for this reduction. Therefore, it can be inferred that CtaA may regulate H_2_S production by modulating the expression of genes involved in H_2_S biosynthesis.

3.3 *ctaA* regulates virulence factors in *S. aureus*

To investigate the role of *ctaA* in virulence regulation, we assessed hemolysin production and staphyloxanthin synthesis *in vitro*, followed by evaluation of virulence *in vivo* using the *G. mellonella* infection model. As shown in Fig. 4A & 4B, under glucose-deficient conditions, the △*ctaA* mutant exhibited a 56% reduction in hemolysin production and a 66% increase in staphyloxanthin production compared to the WT strain. In the presence of 0.25% glucose, the mutant displayed a 43% decrease in hemolysin and a 29% increase in staphyloxanthin. Survival of *G. mellonella* larvae was monitored every 24 h for 5 days (Fig. 4C). The WT group exhibited rapid mortality in the absence of glucose, with an LT₅₀ of 30 h (95% CI: 24–36 h). In contrast, the Δ*ctaA* mutant showed significantly prolonged survival (LT₅₀ > 100 h; \*\**P*< 0.01), confirming that *ctaA* is a critical virulence factor. Interestingly, in the presence of glucose, the survival rate of the Δ*ctaA* group was 57% compared to 65% in the WT group (*P*< 0.05), indicating a glucose-dependent modulation of virulence (Fig. 4D).

**Fig. 4.**
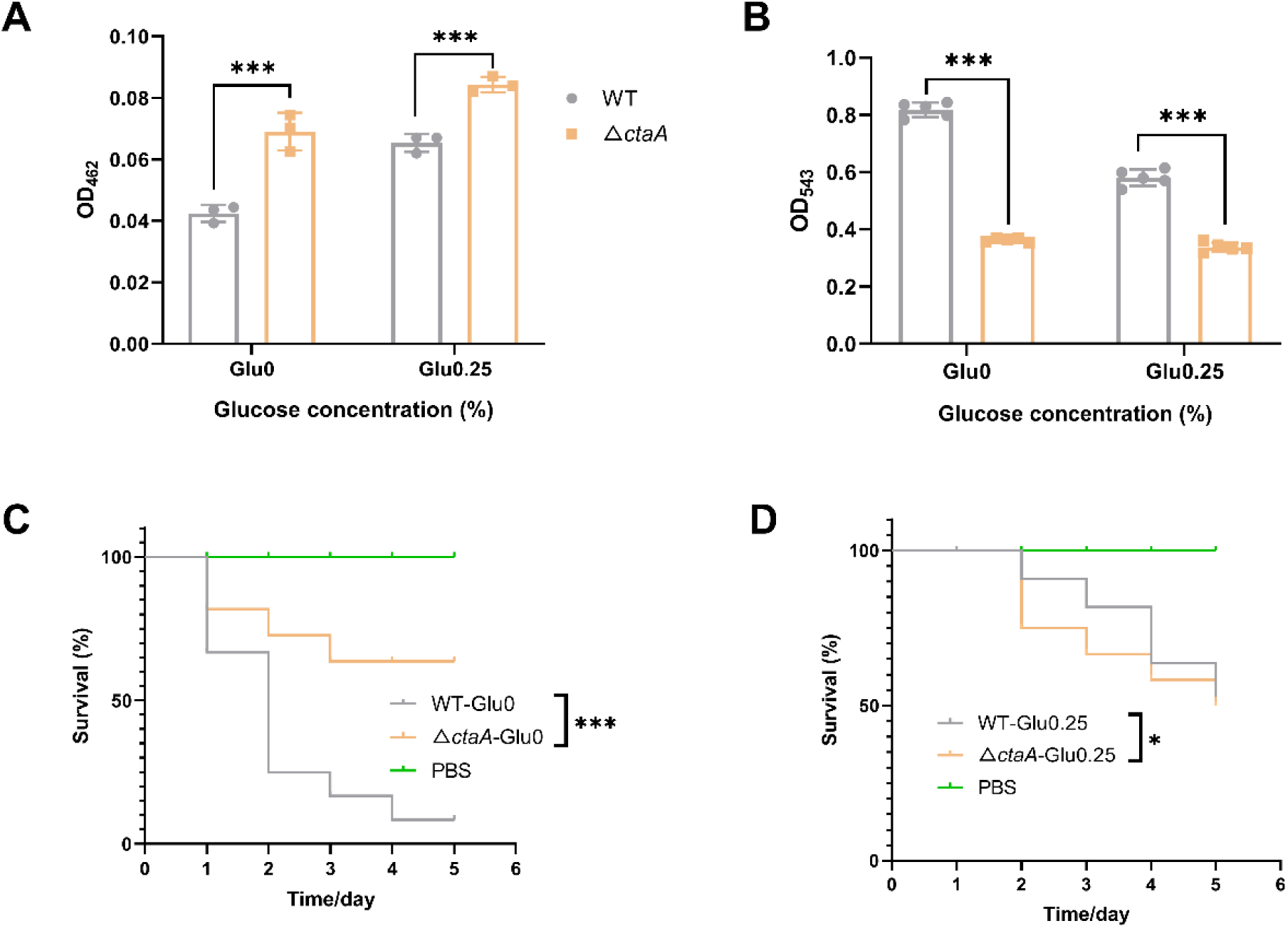
Virulence-related phenotype validation of △*ctaA* mutant. (A) Staphyloxanthin production; (B) Hemolysin formation; (C) *Galleria mellonella* survival rate under glucose-free conditions; (D) *Galleria mellonella* survival rate under 0.25% glucose conditions. Data represents mean±SD from at least n = 3 biological replicates. \*\**P* < 0.01.

3.4 *ctaA* regulates differential antimicrobial resistance in *S. aureus*

In this study, the ability to produce H_2_S was significantly impaired in the △*ctaA* mutant, particularly under glucose-limited conditions. To determine whether the dramatic reduction of H_2_S production in the △*ctaA* mutant affects antimicrobial resistance in *S. aureus*, we measured the MICs of multiple antimicrobial agents against both the WT strain BAA1717 and the △*ctaA* mutant in the presence (0.25% glucose) or absence of glucose.

Glucose availability critically modulated the antimicrobial resistance phenotype mediated by *ctaA*. As shown in Table 2, disruption of *ctaA* led to a moderate increase in antimicrobial resistance under glucose-depleted conditions. However, this effect was markedly more pronounced in glucose-replete environments, where the mutant exhibited a 2– to 4-fold increase in resistance to β-lactams (oxacillin and ampicillin), gentamicin sulfate and erythromycin. Notably, the MIC of daptomycin against the Δ*ctaA* mutant increased two-fold under glucose-depleted conditions, whereas no significant difference was observed in the presence of 0.25% glucose compared to the WT strain.

**Table 2.**
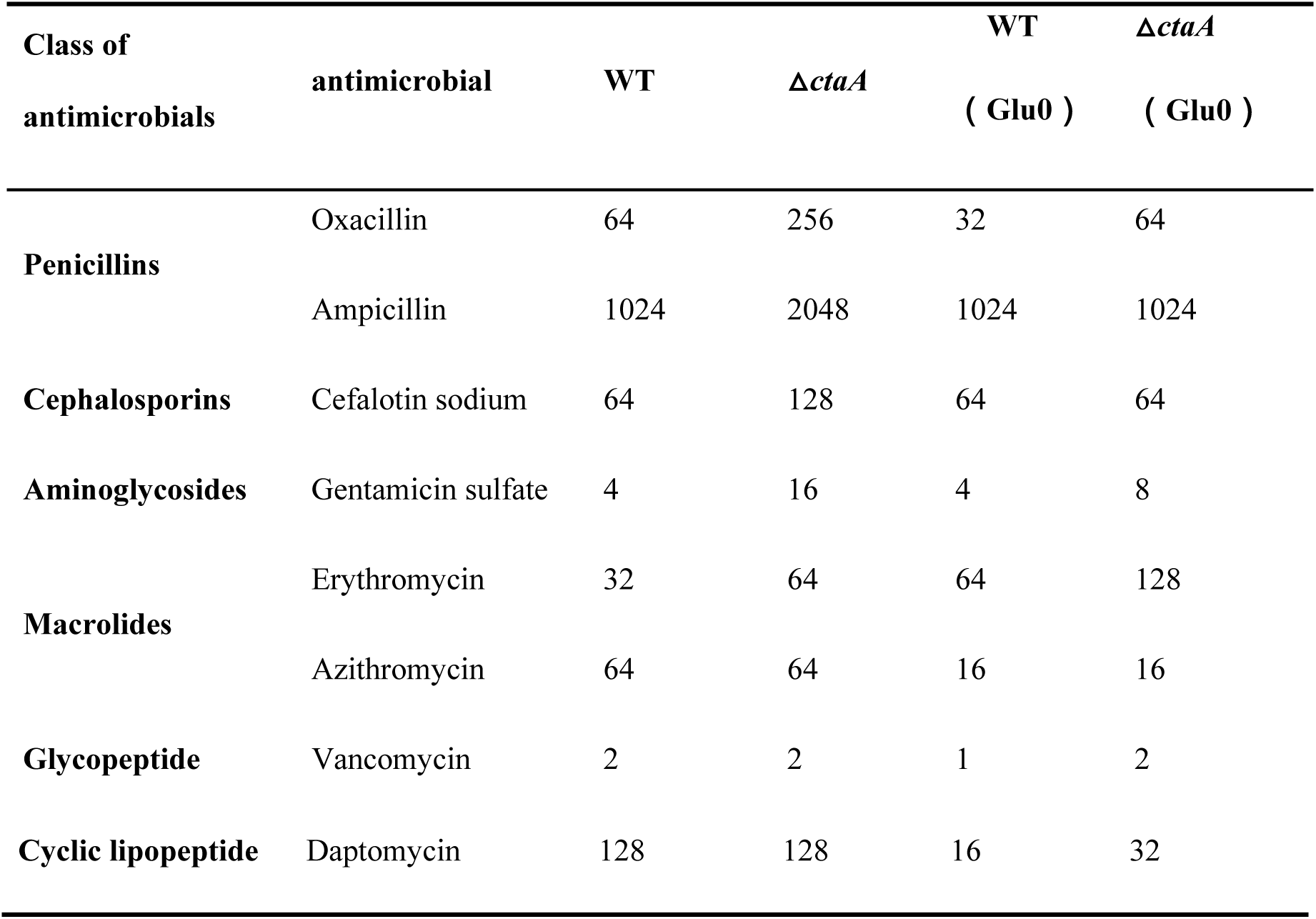
MIC determination (mg/L) of different types of antimicrobial agents against the WT and △*ctaA* strains.

### 3.5 CtaA is required to maintain normal cell morphology

To determine whether deletion of *ctaA* affects cellular morphology, we performed SEM imaging on the Δ*ctaA* mutant and WT strains. As shown in Fig. 4, SEM analysis revealed no significant morphological differences between the mutant and WT cells. Both strains exhibited smooth surfaces and maintained intact cell shapes without evidence of lysis or shrinkage (Fig. 5A). Measurement of maximum cell diameters from SEM images using ImageJ (v1.53) indicated a 12% reduction in the mutant compared to WT cells (Fig. 5B, \*\**P*<0.01).

**Fig. 5.**
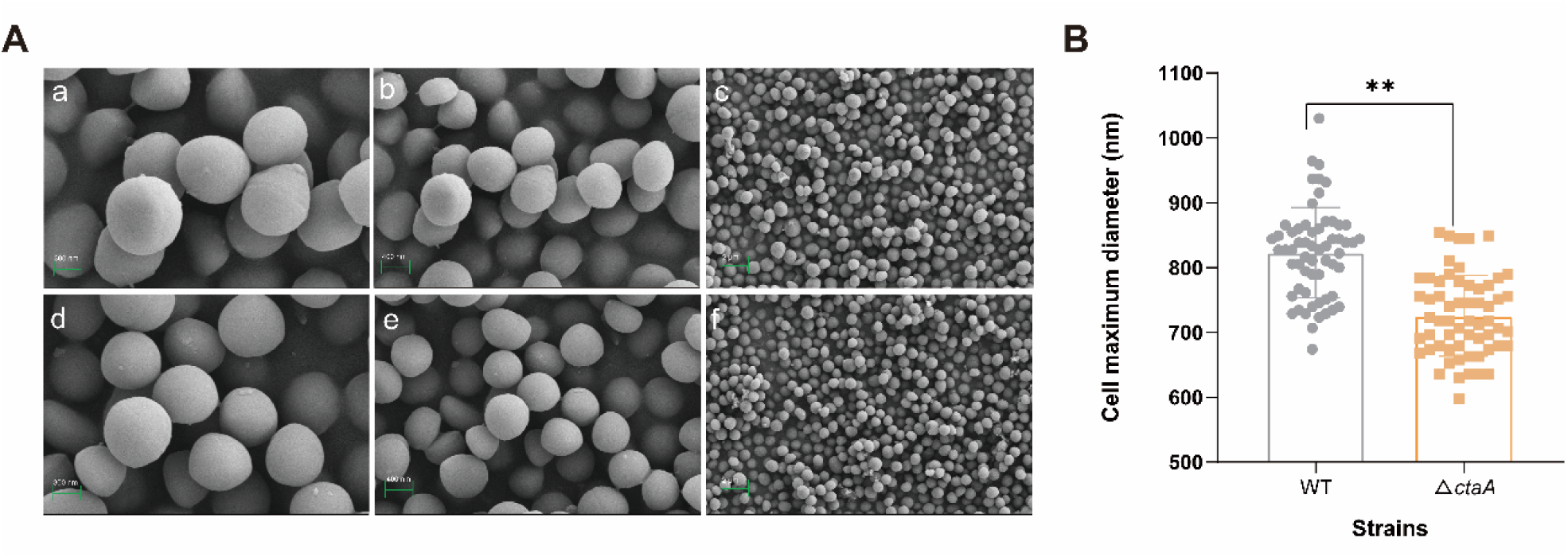
Scanning electron microscopy (SEM) of cell morphology of the WT and △*ctaA* mutant strains. (A) a, b, c: SEM images of the WT cells under different magnification (×30K/20K). d, e, f: SEM images of the △*ctaA* mutant cells at different magnifications (×30K/20K); (B) Maximum Cell diameters quantified as the average of 60 random selected cells by Image J. \*\**P* < 0.01.

TEM analysis showed that under glucose-supplemented conditions, both WT and △*ctaA* strains displayed significantly thicker and more compact cell walls compared to glucose-depleted controls. Quantitative TEM analysis further demonstrated that *ctaA* deletion consistently increased cell wall thickness regardless of glucose availability (Fig. 6A). In glucose-depleted medium, the cell walls of the Δ*ctaA* mutant were 21% thicker than those of the WT strain (Fig. 6B, 30.7 nm vs. 25.2 nm; *P* < 0.01). Although supplementation with 0.25% glucose led to cell wall thickening in both strains, the Δ*ctaA* mutant still exhibited significantly thicker walls (Fig. 6B, 36.1 nm vs. 31.8 nm for WT, *P* < 0.01). Additionally, when cultured on TSA plates, the Δ*ctaA* mutant formed smaller colonies with more intense pigmentation compared to the WT strain (Fig. S1), consistent with our SEM observations and staphyloxanthin quantitation data (Fig. 4A & Fig. 5).

**Fig. 6.**
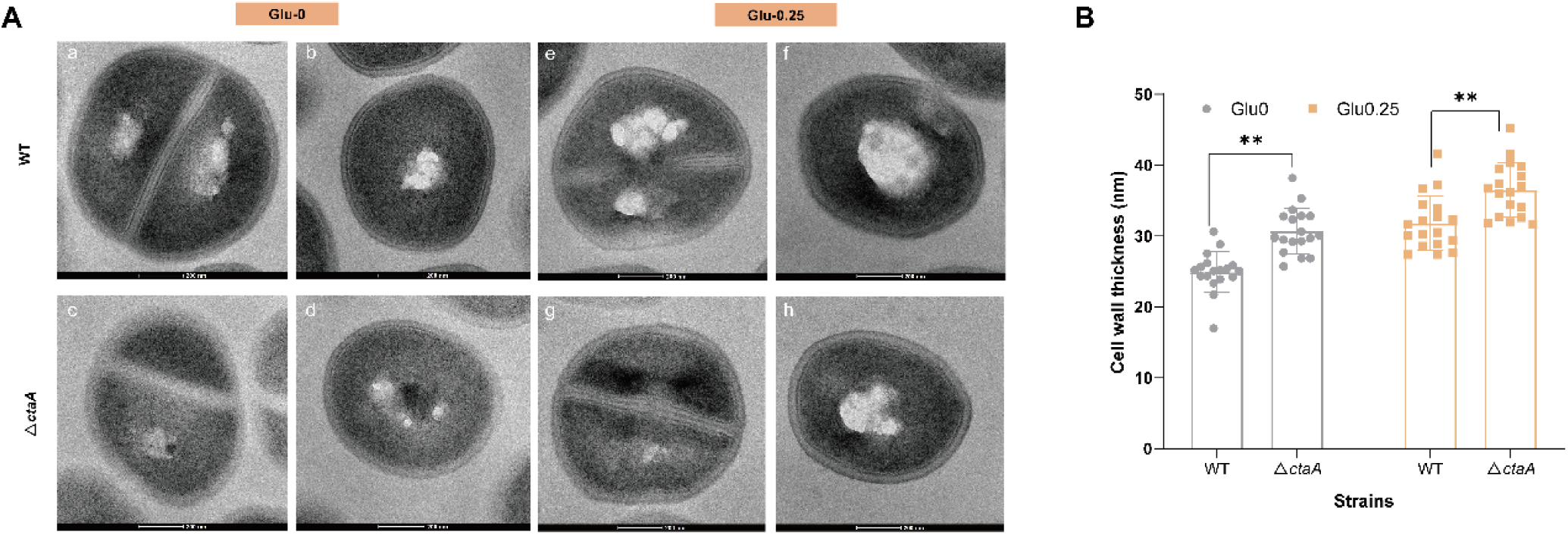
Representative TEM images exhibiting cell morphology of the WT and *ctaA* mutant strains in the absence and presence of glucose. (A) a, b: TEM images of two representative WT cells in the absence of glucose (×55K). c, d: TEM images of two representative △*ctaA* cells in the absence of glucose (×55K); e, f: TEM images of two representative WT cells in the presence of glucose (×55K); g, h: TEM images of two representative △*ctaA* cells in the presence of glucose (×55K). (B) Cell thickness quantified as the average of 18 random selected cells by Image J. \*\**P* < 0.01.

3.6 Deletion of *ctaA* alters the transcriptome profiles of *S. aureus*

To elucidate the role of CtaA in orchestrating transcriptomes associated with H_2_S biosynthesis, we performed RNA-seq analysis of the WT and isogenic Δ*ctaA* strains cultured in glucose-replete (0.25%) and glucose-free media. Comparative transcriptomic analysis revealed a total of 199 upregulated and 110 downregulated DEGs under glucose-free conditions, whereas 312 upregulated and 195 downregulated DEGs were identified under glucose-replete conditions (Fig. 7A&7B).

**Fig. 7.**
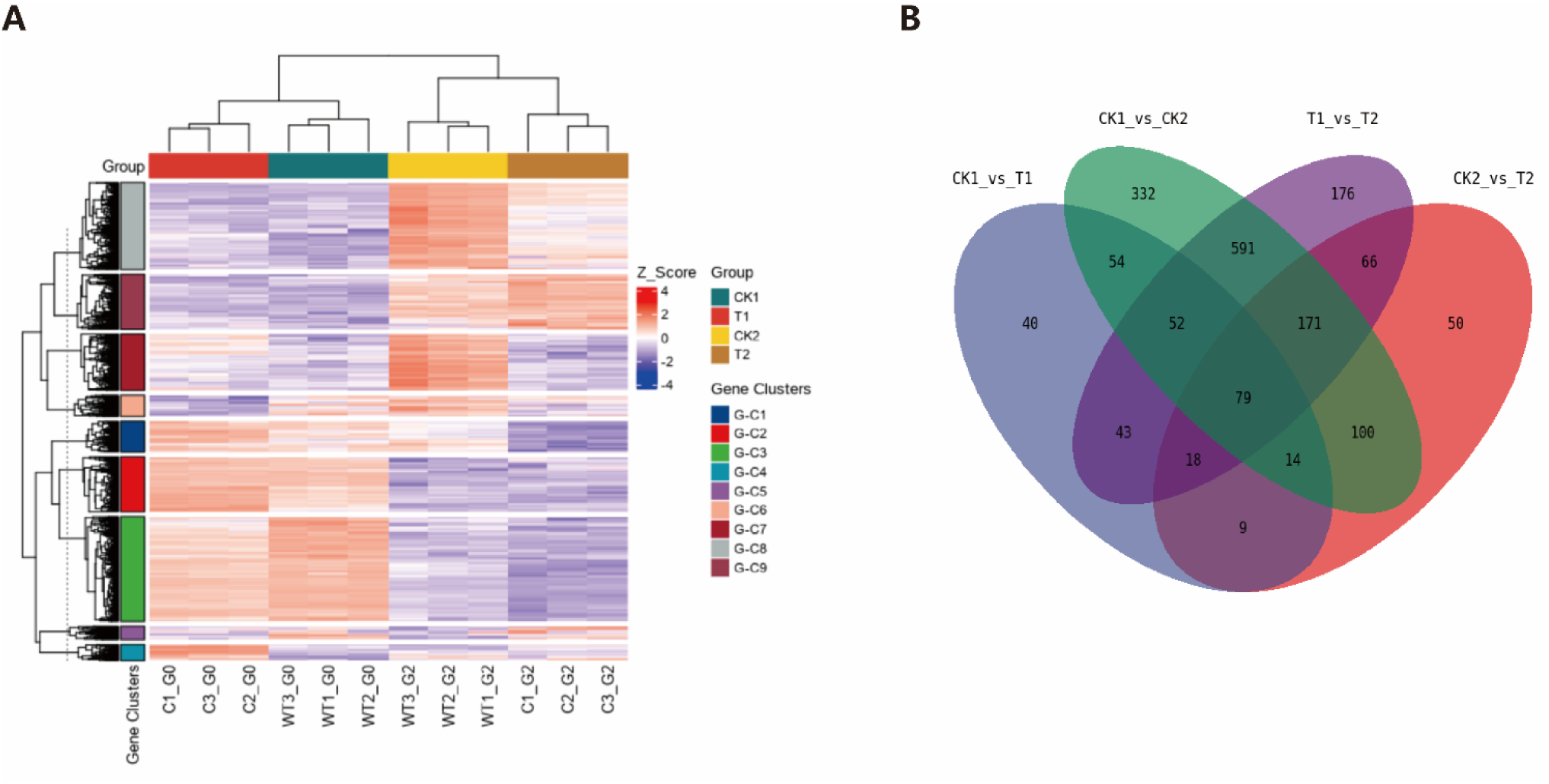
Transcriptomic analysis of Staphylococcus aureus wild-type (WT) and Δ*ctaA* strains under glucose-limited (Glu0) and glucose-replete (Glu0.25) conditions. (A) Hierarchical clustering heatmap of differentially expressed genes (C1-G0/C2-G0/C3-G0: biological replicates of Δ*ctaA* under Glu0; WT1-G0/WT2-G0/WT3-G0: biological replicates of WT under Glu0; C1-G2/C2-G2/C3-G2: biological replicates of Δ*ctaA* under Glu0.25; WT1-G2/WT2-G2/WT3-G2: biological replicates of WT under Glu0.25); Genes are divided into 9 expression-based clusters (G-C1 to G-C9), as indicated by the colored bars on the left. Color intensity represents the Z-score normalized expression level (red: high expression; blue: low expression). (B) Venn diagram showing the overlap of differentially expressed genes across four comparisons: CK1_vs_T1 (WT vs. Δ*ctaA* at Glu0), CK2_vs_T2 (WT vs. Δ*ctaA* at Glu0.25), CK1_vs_CK2 (WT at Glu0 vs. Glu0.25), and T1_vs_T2 (Δ*ctaA* at Glu0 vs. Glu0.25). Data represents mean±SD from n = 3 biological replicates.

To further understand the metabolic pathways affected by *ctaA*, the identified DEGs were categorized into Gene Ontology (GO) functional groups. Under glucose-free conditions, 309 DEGs were associated with the arginine metabolic process, alpha-amino acid metabolic process, and active ion transmembrane transporter activity process (Fig. 8A). In contrast, glucose-supplementation led to a significant increase in DEGs linked to the organonitrogen compound, carboxylic acid metabolic process, oxoacid metabolic process, and intracellular anatomical structures–– pathways not prominently enriched under glucose-free conditions (Fig. 8B). Collectively, these DEGs could be grouped into three major regulatory categories: (i) arginine metabolic process, (ii) biosynthesis of glutamine family amino acid, and (iii) active ion transmembrane transporter activity.

**Fig. 8.**
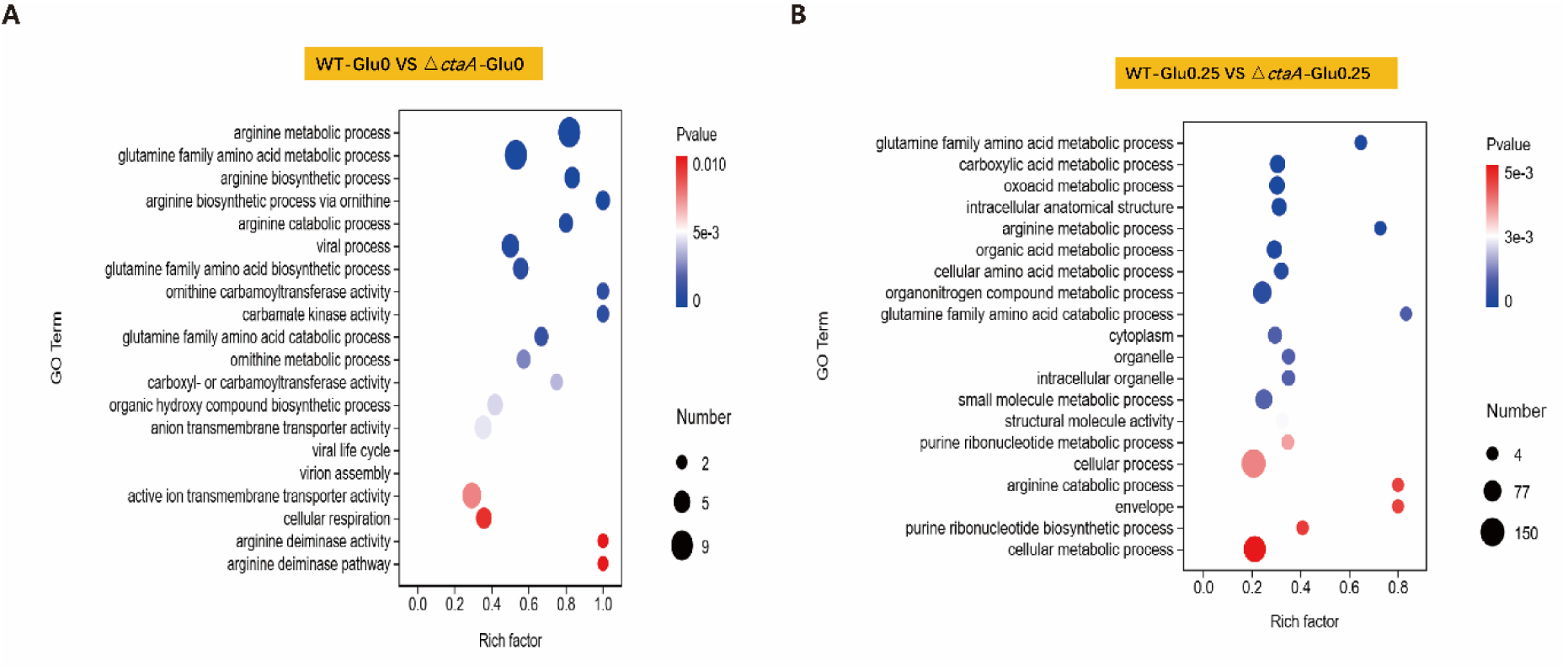
Gene Ontology (GO) enrichment analysis of differentially expressed genes in Staphylococcus aureus wild-type (WT) and Δ*ctaA* strains under glucose-limited (Glu0) and glucose-replete (Glu0.25) conditions. (A) WT-Glu0 vs. Δ*ctaA*-Glu0; (B) WT-Glu0.25 vs. Δ*ctaA*-Glu0.25. Bubble size represents the number of enriched genes, and color indicates statistical significance (*P*-value). Data represents mean±SD from n = 3 biological replicates.

### 3.7 *ctaA* regulates H_2_S production-related genes and influences L-cysteine transport

To investigate how *ctaA* modulates endogenous H_2_S production in *S. aureus*, we analyzed the expression of genes involved in H_2_S biosynthesis using RNA-seq data. As shown in Fig. 9A, significant changes (ranging from 1.01– to 5.86-fold) were observed in key genes associated with cysteine metabolism, including the transcriptional regulator (*cymR*), the cysteine transporter (*tcyA*), and cystathionine β-lyase (*metC*). These findings suggest that the deletion of *ctaA* disrupts both cysteine uptake and the expression of genes involved in endogenous H_2_S generation.

**Fig. 9.**
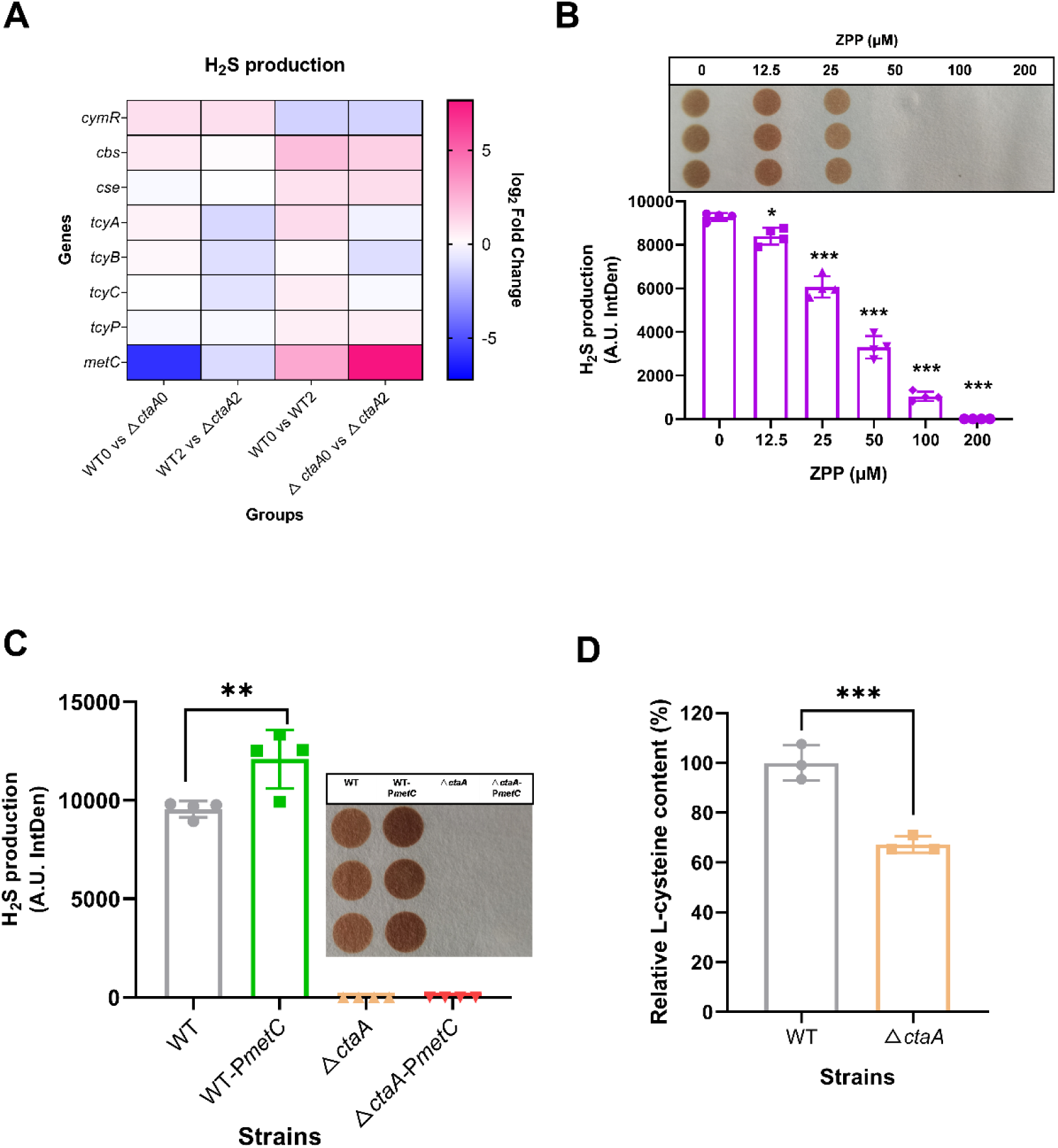
*ctaA* regulates the genes involved in H_2_S production and influences L-cysteine transport. (A) Heatmap showing relative transcription levels of H_2_S production-related genes (WT0: wild-type without glucose; Δ*ctaA*0: Δ*ctaA* mutant without glucose; WT2: wild-type with 0.25% glucose; Δ*ctaA*2: Δ*ctaA* mutant with 0.25% glucose); (B) Inhibitory effect of the cytochrome oxidase inhibitor ZPP (0–200 μM) on H_2_S production in the wild-type strain; (C) Effect of *metC* overexpression on H_2_S production in wild-type (WT) and Δ*ctaA* mutant strains; (D) Intracellular Cys content in wild-type and Δ*ctaA* mutant strains. **, *P* < 0.01; ***, *P* < 0.001. Data represents mean±SD from at least n = 3 biological replicates.

Given that *ctaA* encodes heme A synthase—a critical component of the bacterial electron transport chain—we hypothesized that its loss may impair respiratory function, thereby affecting H_2_S production. To test this, we treated the WT strain with ZPP, a known inhibitor of the bacterial electron transport chain. As illustrated in Fig. 9B, even at a low concentration (25 μM), ZPP significantly suppressed H_2_S production, with complete inhibition observed at 50 μM. Furthermore, we examined whether iron supplementation could rescue H_2_S production in the Δ*ctaA* mutant by restoring heme biosynthesis. However, FeCl_3_ supplementation failed to restore H_2_S production in the Δ*ctaA* mutant (Fig. S2), indicating *ctaA* does not regulate H_2_S production through iron homeostasis.

Among sulfur metabolic genes, *metC*, which encodes cystathionine β-lyase responsible for cleaving cystathionine into homocysteine, α-ketobutyrate, and ammonia, exhibited the most significant downregulation (5.86-fold). While bacteria typically cannot directly use homocysteine as a primary substrate for H_2_S synthesis, homocysteine can indirectly contribute to H_2_S production by serving as a sulfur source through interconnected metabolic pathways. Given its potential role as an intermediate for H_2_S biosynthesis, we overexpressed *metC* in the △*ctaA* mutant; however, this did not restore H_2_S production. In contrast, *metC* overexpression in the WT strain significantly enhanced H_2_S production (Fig. 9C). RNA-seq analysis revealed substantial alterations in genes involved in amino acid biosynthesis, transport, and phosphate uptake. Consistent with these findings, intracellular cysteine levels were reduced by approximately 40% in the △*ctaA* mutant strain (Fig. 9D).

### 3.8 CtaA regulates cysteine uptake in *S. aureus* by modulating the activity of the L-cysteine importer TcyP

Exogenous cysteine acquisition serves as the primary substrate source for endogenous H_2_S production in *S. aureus*. RNA-seq analysis revealed that the deletion of *ctaA* resulted in a significant upregulation of *cymR*, the major transcriptional repressor of cysteine metabolism (Fig. 9A). Consistent with this, RT-qPCR analysis showed marked downregulation of genes involved in L-cysteine transport (*tcyP*) and H_2_S production (*mccB*) (Fig. 10A). To further assess whether *ctaA* affects the transcriptional regulation of sulfur metabolism-related genes, we performed a GFP reporter fusion assay. As shown in Fig. 10B, promoter activities of *cymR, cse,* and *tcyP* were significantly altered in the Δ*ctaA* mutant, with *tcyP* promoter activity reduced by more than 75%. Given that TcyP is the principal L-cysteine transporter in *S. aureus*, we next examined whethe*r ctaA* affects TcyP expression at the protein level. Using a His-tag fusion system, we quantified TcyP protein levels in the WT and Δ*ctaA* strains. Fig. 10C shows that TcyP expression decreased by over 40% in the mutant, consistent with the reduced intracellular L-cysteine levels observed in the Δ*ctaA* strain (Fig. 10D). Moreover, we overexpressed *tcyP* in both the WT and Δ*ctaA* mutant strains. As illustrated in Fig. 10E, complementation fully restored the severe reduction in endogenous H_2_S production observed in the Δ*ctaA* mutant under glucose-depleted conditions. In contrast, *tcyP* overexpression had no significant effect on endogenous H_2_S levels in the WT strain (Fig. 10F). Collectively, these findings demonstrate that *ctaA* modulates endogenous H_2_S production in *S. aureus* by regulating the expression and function of the L-cysteine transporter TcyP.

**Fig. 10.**
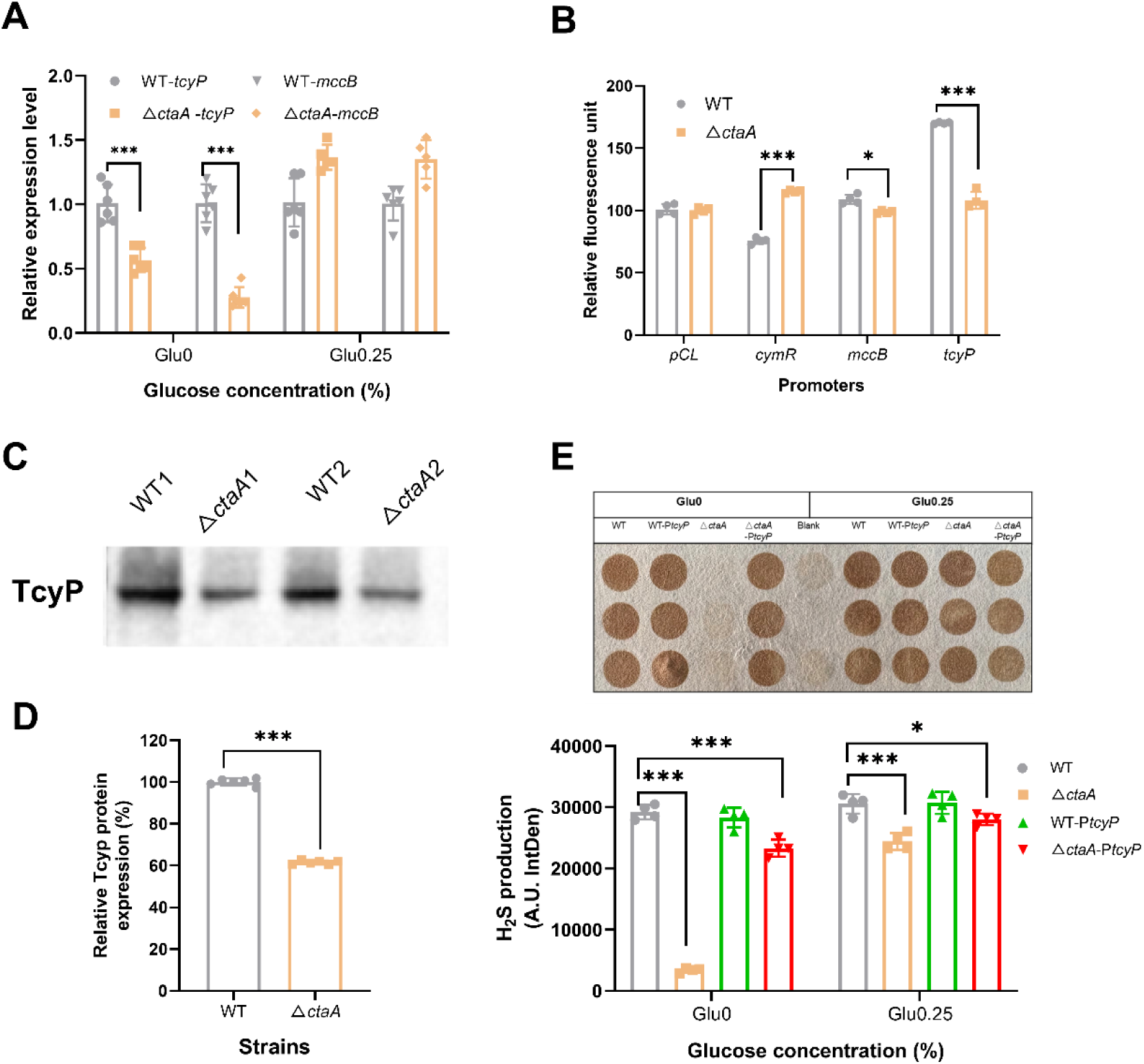
CtaA regulates cysteine uptake in *S. aureus* by modulating the activity of the L-cysteine importer TcyP. (A) Relative transcription levels of key genes involved in H_2_S production; (B) Effect of *ctaA* on the promoter activity of H_2_S production-related genes (detected via GFP reporter assay); (C) TcyP protein expression levels determined by Western blot; (D) Relative quantification of TcyP protein expression in wild-type (WT) and Δ*ctaA* strains; (E) Relative quantification of H_2_S production in WT and ΔctaA strains overexpressing *tcyP* under glucose-limited and glucose-replete conditions. *, *P* < 0.05; ***, *P* < 0.001. All assays were performed in triplicate.

### 3.9 CtaA regulates H_2_S production by modulating arginine metabolism

Under glucose-depleted or energy-limiting conditions, arginine serves as a critical alternative energy source for *S. aureus.* RNA-seq analysis revealed that deletion of *ctaA* resulted in significant upregulation of genes involved in arginine metabolism, particularly in the absence of glucose, with fold changes ranging from 1.16 to 3.09 (Fig. 11A). This suggests that the *ctaA* mutant strain relies on enhanced arginine uptake and catabolism to maintain growth under metabolic stress. Notably, the phosphate ABC transporter genes *pstSCAB* and their regulator *phoU* were downregulated by 1.03– to 2.28-fold in the △*ctaA* mutant under glucose restriction (Fig. 11B).

**Fig. 11.**
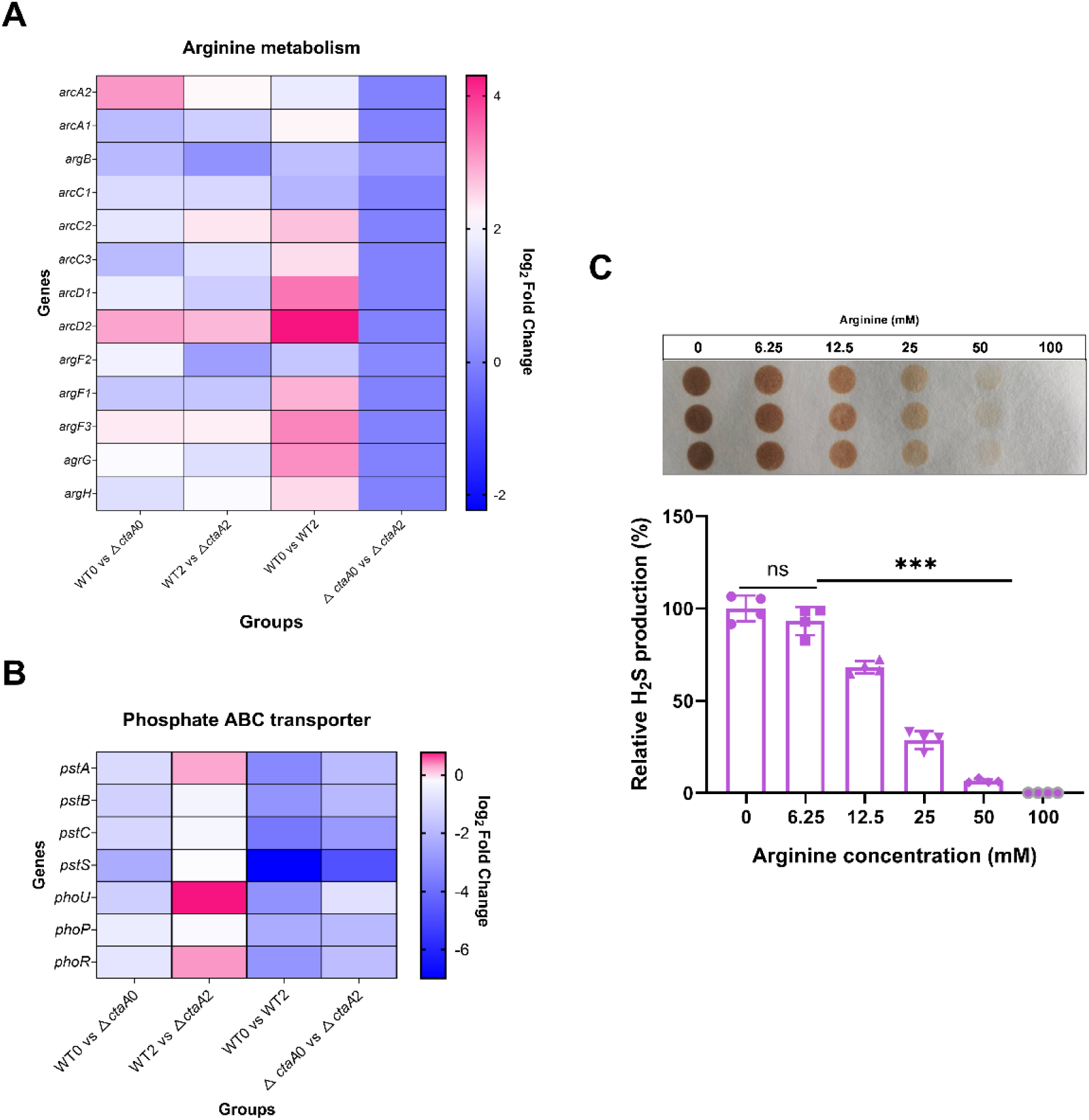
Deletion of *ctaA* reprograms arginine metabolism in a glucose-responsive manner to regulate H_2_S production. (A) Heatmap showing relative transcription levels of arginine metabolism-related genes (WT0: wild-type without glucose; Δ*ctaA*0: Δ*ctaA* mutant without glucose; WT2: wild-type with 0.25% glucose; Δ*ctaA*2: Δ*ctaA* mutant with 0.25% glucose); (B) Heatmap showing relative transcription levels of phosphate transporter-related genes; (C) H_2_S production in *S. aureus* BAA1717 under normal conditions and media with different concentrations of arginine, measured by lead acetate test paper and quantified using Image J software. ***, *P* < 0.001. All assays were performed in triplicate.

To investigate whether increased arginine availability affects endogenous H_2_S production, we supplemented the culture medium with exogenous arginine at varying concentrations and measured H_2_S production levels. As shown in Fig. 11C, even a low concentration of arginine (6.25 mM) significantly suppressed H_2_S production. This inhibition exhibited a dose-dependent pattern, with higher arginine concentrations leading to progressively greater reductions in H_2_S levels. At 50 mM arginine, H_2_S production was nearly completely abolished. These findings indicate that intracellular arginine levels play a crucial role in modulating endogenous H_2_S generation in *S. aureus*.

### 3.10 CtaA regulates H_2_S production by modulating NO generation

NO is another key gaseous signaling molecule and arginine serves as a primary precursor for endogenous NO biosynthesis in bacteria. Our RNA-seq analysis revealed that in the △*ctaA* mutant strain, genes involved in arginine metabolism were upregulated, along withthose associated with NO production and NO resistance--particularly under glucose-deprived conditions. As shown in Fig. 12A, in the absence of glucose, significant upregulation (ranging from 1.41– to 1.93-fold) was observed in genes encoding nitric oxide synthase (nos), L-lactate dehydrogenases (*ldh1, ldh2*), flavohemoprotein (*hmp*), nitrite reductase components *(nirB, nirD*), and subunits of respiratory nitrate reductase (*narG, narH, narJ*).

**Fig. 12.**
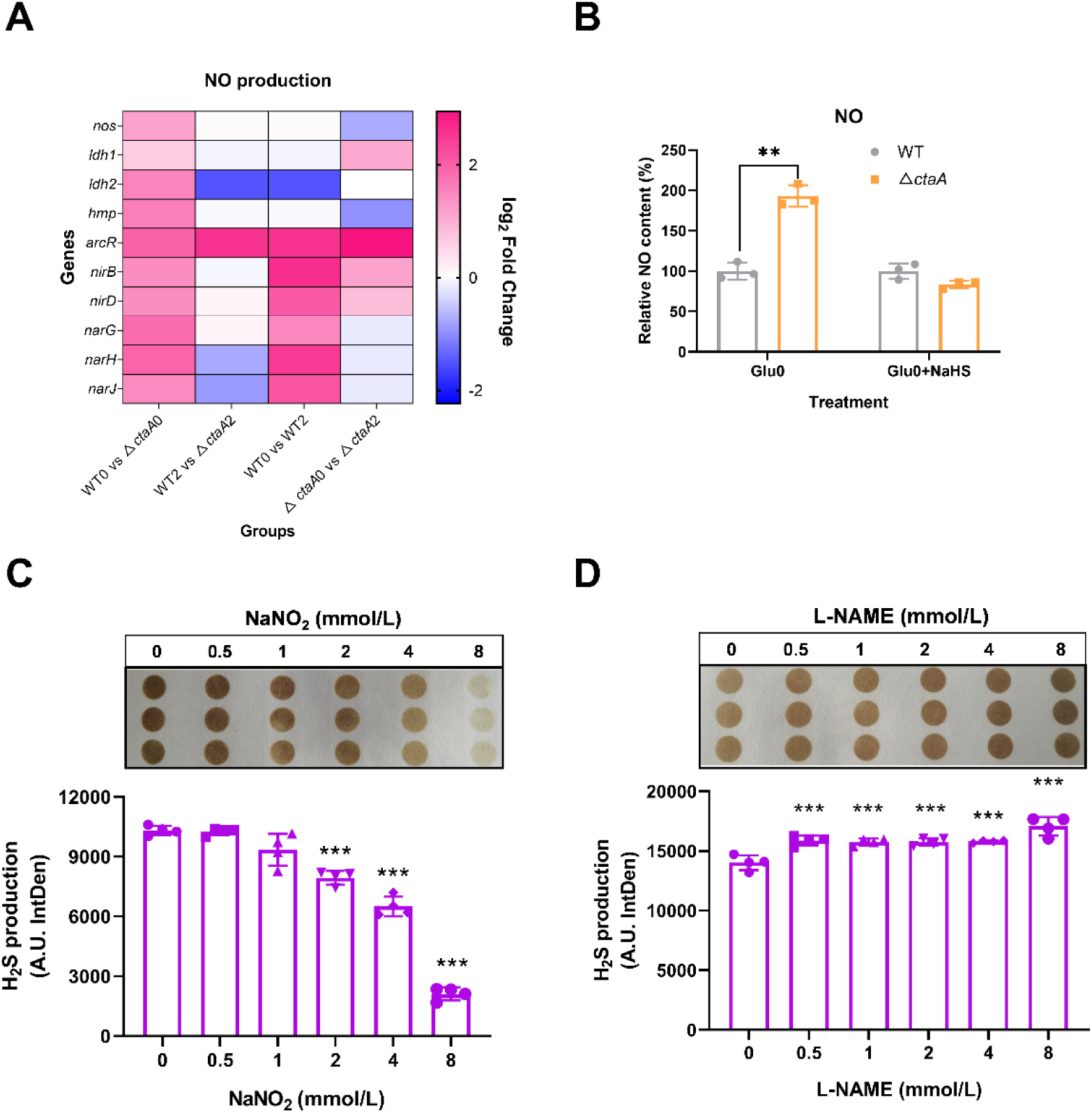
*ctaA* influences H_2_S production through NO regulation under glucose-limited conditions. (A) Heatmap analysis of relative transcription levels of NO production-related genes (WT0: wild-type without glucose; Δ*ctaA*0: Δ*ctaA* mutant without glucose; WT2: wild-type with 0.25% glucose; Δ*ctaA*2: Δ*ctaA* mutant with 0.25% glucose); (B) NO content with or without NaHS supplementation under glucose-free conditions; (C) Dose-dependent effect of the NO donor NaNO₂ (0–8 mM) on H_2_S production in *S. aureus* BAA1717; (D) Dose-dependent effect of the NO inhibitor L-NAME (0–8 mM) on H_2_S production in *S. aureus* BAA1717. **, *P* < 0.01; ***, *P* < 0.001. All assays were performed in triplicate.

Consistent with these transcriptomic findings, quantitative measurement of intracellular NO levels demonstrated an 80% increase in the △*ctaA* mutant under glucose-limiting conditions, while no significant change was observed under glucose-replete conditions (Fig. 12B). This further supports the notion that CtaA modulates NO production in a carbon source-dependent manner.

Next, we investigated whether exogenous H_2_S, delivered via the donor NaHS, affects endogenous NO production in the △*ctaA* mutant under glucose deprivation. Our results showed that NaHS supplementation significantly suppressed NO generation in the mutant strain.

To further explore the interplay between NO and endogenous H_2_S in *S. aureus*, we applied the NO donor sodium nitrite (NaNO₂) and assessed its impact on H_2_S production. As illustrated in Fig. 12C, 1 mM NaNO₂ markedly suppressed H_2_S synthesis, with near-complete inhibition observed at 8 mM. Conversely, treatment with the NO synthase inhibitor L-NAME significantly enhanced endogenous H_2_S production (Fig. 12D). These findings collectively support the notion that NO negatively regulates H_2_S biosynthesis in *S. aureus*.

## 4. Discussion

The ability of *S. aureus* to withstand host-imposed oxidative stress is a cornerstone of its pathogenicity[35], with endogenous H_2_S serving as a critical component of this redox shield[36]. While the enzymatic source of H_2_S (CSE) and its cytoprotective roles are well established[1], the upstream regulatory networks that dynamically tune H_2_S production in response to metabolic cues remain poorly defined. In this study, we uncover a previously unrecognized metabolic-redox axis wherein the respiratory chain integrity, governed by CtaA, acts as a master switch controlling H_2_S homeostasis. Our findings reveal that under glucose starvation—a hallmark of the host infection niche—CtaA deficiency triggers a profound metabolic reprogramming that collapses H_2_S production while simultaneously inducing a compensatory NO surge. This work redefines CtaA not merely as a respiratory assembly factor, but as a central conductor orchestrating the crosstalk between bioenergetics, sulfur metabolism, and gasotransmitter signaling.

In this study, *ctaA* deletion markedly increased staphyloxanthin production in *S. aureus*. As reported previously, loss of *ctaA* impairs aerobic respiration and elevates intracellular ROS, which activates σB-dependent transcription of the *crtOPQMN* operon and thus causes higher staphyloxanthin [37]. Meanwhile, hemolysin synthesis was dramatically reduced in the Δ*ctaA* mutant, consistent with prior findings[37]. Similar phenotypic changes were also observed in the *ctaB*-deficient strain, confirming that these alterations result from impaired respiratory function[20]. Notably, glucose concentration did not affect the trends of these two phenotypes, which differs from the regulatory pattern of endogenous H_2_S and indicates that endogenous H_2_S and respiratory dysfunction modulate these virulence traits through distinct pathways.

Traditionally, H_2_S biosynthesis in bacteria has been viewed through the lens of substrate availability and canonical sulfur regulators like CymR [38, 39]. However, our data suggest a more complex, top-down regulation driven by the functional status of electron transport chain (ETC). The near-abolition of H_2_S in the △*ctaA* mutant under glucose limitation cannot be solely attributed to a passive energy deficit. Instead, we propose a model of retrograde signaling: the failure to assemble the cytochrome *aa_3_* oxidase (due to lack of heme A) generates a specific metabolic signal that actively represses the cysteine uptake machinery (TcyP).

Our transcriptomic analyses and Western Blot results pinpoint the L-cysteine transporter TcyP [40] as the critical effector of this regulation. The downregulation of *tcyP*, mediated by the upregulation of the repressor *cymR*, creates a “substrate bottleneck” that starves CSE of its precursor. This mechanism likely serves as an adaptive strategy to prevent the accumulation of toxic sulfur intermediates when the ETC is compromised, or to redirect limited carbon flux away from non-essential antioxidant synthesis toward immediate survival pathways [41]. More importantly, the overexpression of *tcyP* in △ *ctaA* strain could completely rescue its H_2_S production under glucose deficient conditions, confirming *ctaA* regulates the H_2_S production via cysteine metabolism inhibition. These results are consistent with a previous study by Timmie et al. on *Fusobacterium nucleatum*, which demonstrated that inactivation of the respiratory enzyme complex Rnf abolished endogenous H_2_S production from cysteine, thereby linking disrupted energy metabolism to impaired cysteine catabolism and H_2_S synthesis [42]. These findings also aligns with emerging concepts in mitochondrial biology where respiratory dysfunction retrograde-regulates nuclear gene expression [43, 44]; here, we demonstrate an analogous bacterial communication where respiratory status dictates cytoplasmic metabolic flux.

The transcriptomic landscape of *S. aureus* undergoes a dramatic shift upon *ctaA* deletion. Facing an energy crisis induced by impaired aerobic respiration and glucose scarcity, the bacterium prioritizes the Arginine Deiminase (ADI) pathway over cysteine metabolism. The ADI pathway, capable of generating ATP via substrate-level phosphorylation independent of the ETC and oxygen, represents a vital “lifeboat” mechanism [45, 46].

Our observation that arginine accumulation correlates with H_2_S suppression suggests a metabolic trade-off. By upregulating the ADI pathway and downregulating cysteine uptake, *S. aureus* sacrifices its long-term oxidative defense (H_2_S) to secure immediate energy currency (ATP) and pH homeostasis. This rewiring explains the paradoxical phenotype of the △*ctaA* mutant: while it becomes hypersensitive to exogenous oxidative stress due to H_2_S depletion, it exhibits altered resistance profiles potentially linked to the metabolic state shifts or cell wall modifications associated with arginine metabolism. This highlights the plasticity of bacterial metabolism[47], where the hierarchy of nutrient utilization is dynamically recalibrated by the integrity of the respiratory chain [48].

Perhaps the most striking finding of this study is the reciprocal regulation between H_2_S and NO in *S. aureus*. We observed that the collapse of H_2_S in the Δ *ctaA* mutant is accompanied by a significant surge in endogenous NO, driven by the upregulation of NO-producing genes and the accumulation of arginine (the NO precursor). Functional assays confirmed a possible antagonistic relationship: exogenous H_2_S suppresses NO, while NO donors inhibit H_2_S production. This H_2_S-NO crosstalk mirrors mechanisms described in mammalian systems, where these gasotransmitters compete for heme targets or modulate each other’s synthesis to fine-tune vascular tone and oxidative stress responses [49, 50]. Previous studies have documented interactions between gaseous signaling molecules in both eukaryotes and prokaryotes [51–53]. For instance, in *E.coli*, NO production is elevated in H_2_S-deficient cells, while H_2_S production increases in NO-deficient cells compared to WT strains [4]. In *S. aureus*, we hypothesize that this interaction serves as a compensatory redox switch. When the primary antioxidant (H_2_S) is unavailable due to metabolic constraints, the bacterium may upregulate NO to modulate respiration (via inhibition of residual oxidases) or to activate alternative stress response pathways (e.g., via S-nitrosylation of key enzymes). However, our data suggest that this compensatory NO surge is insufficient to fully restore virulence or oxidative tolerance, underscoring the non-redundant, essential role of H_2_S in protecting *S. aureus* during host invasion. The disruption of this delicate gas balance by *ctaA* deletion may leave the pathogen vulnerable to host immune clearance.

The relevance of our findings extends directly to the pathogenesis of *S. aureus* infections. Within abscesses, biofilms, and phagosomes, bacteria face the dual challenge of nutrient deprivation (specifically glucose) [18] and intense oxidative attack from immune cells [54, 55]. Our study demonstrates that CtaA is indispensable for maintaining the H_2_S-mediated redox shield under these exact conditions. The attenuation of virulence in the *ctaA* mutant in the *G. mellonella* model, especially under glucose starved condition, likely stems from the inability to sustain H_2_S levels when glucose is scarce, rendering the bacteria susceptible to neutrophil-derived ROS.

From a therapeutic perspective, targeting CtaA offers a distinct advantage over direct CSE inhibition. Since CSE is highly conserved between bacteria and humans, direct inhibitors risk host toxicity. In contrast, CtaA-mediated regulation of H_2_S appears to be a bacteria-specific adaptation linking respiration to virulence. Disrupting the CtaA-H_2_S axis could effectively “disarm” the bacterial redox defense system specifically within the hostile host environment, sensitizing MRSA to both host immunity and conventional antibiotics. This strategy exploits the pathogen’s own metabolic dependency, turning its adaptation mechanism into an Achilles’ heel.

## 5. Conclusion

In conclusion, this study elucidates a novel regulatory circuit in *S. aureus* where heme A biosynthesis governs endogenous H_2_S production through a complex interplay of respiratory integrity, metabolic reprogramming, and gasotransmitter crosstalk (illustrated in Fig. 13). We propose that CtaA functions as a metabolic-redox hub, ensuring that H_2_S synthesis is tightly coupled to the bacterium’s bioenergetic capacity and environmental context. These insights not only advance our understanding of bacterial redox biology but also highlight the CtaA-dependent metabolic network as a promising target for next-generation antivirulence therapies. Future studies utilizing *in vivo* models and redox proteomics will be crucial to map the specific protein targets of H_2_S and NO (e.g., S-sulfhydration vs. S-nitrosylation) within this regulatory framework, further unraveling the sophistication of bacterial gas signaling.

**Fig. 13.**
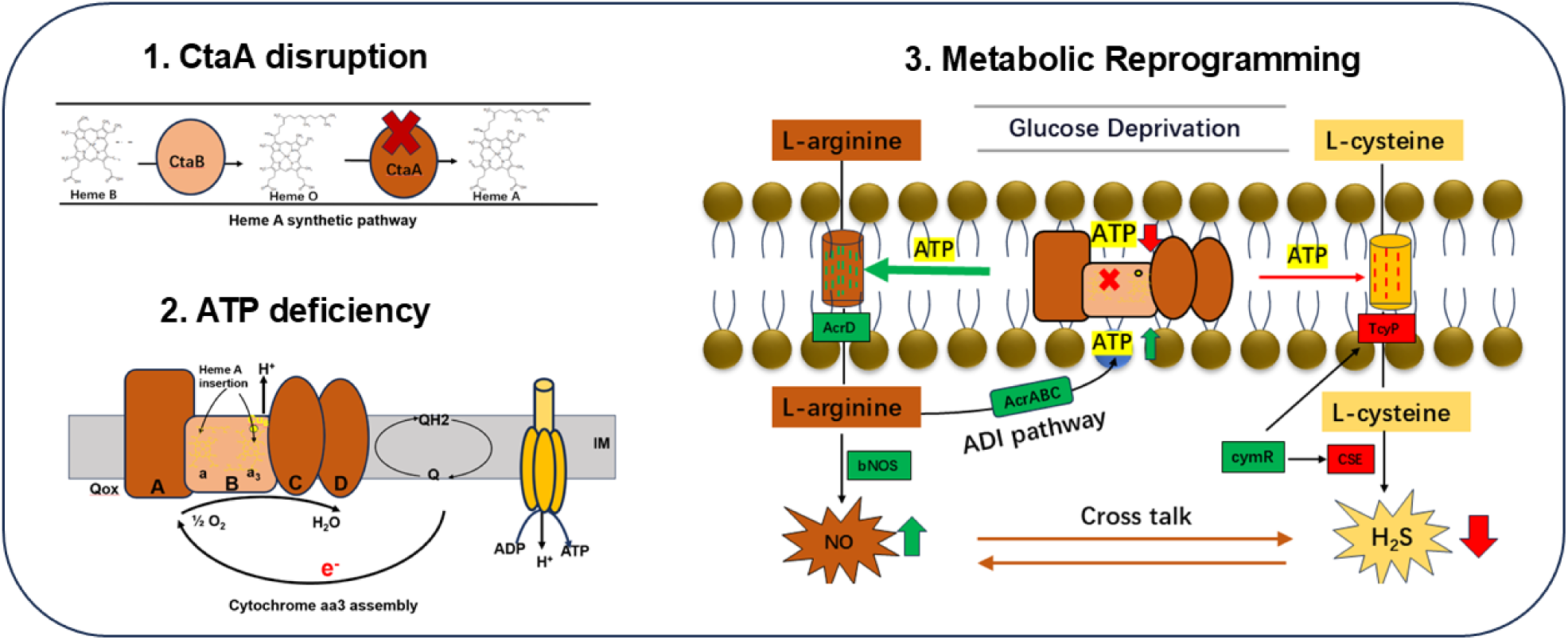
Proposed model for the regulation of endogenous H_2_S production, ATP level, and arginine metabolism by the *ctaA* gene. (1) Deletion of *ctaA* disrupts the conversion of heme O to heme A; (2) Impaired heme A synthesis results in defective assembly of cytochrome oxidase *aa3*, thereby reducing ATP production through this pathway; (3) Reduced ATP output triggers metabolic reprogramming, downregulating secondary metabolic processes such as endogenous H_2_S synthesis, while activating compensatory ATP-generating pathways, including arginine metabolism, to sustain essential cellular functions. Genes upregulated and downregulated in △*ctaA* mutant compared to the WT are highlighted in green and red, respectively.

## CRediT authorship contribution statement

Jiahui Li: Conceptualization, Methodology, Investigation, Formal analysis, Writing – review & editing, Writing – original draft. Mu He & Wanwan Hou: Investigation, Writing – review & editing. Zengfeng Zhang & Zifeng Mai: Investigation. Xiaorong Tian, Nan Zhong & Shimo Kang: Methodology, Investigation. Chunlei Shi: Conceptualization, Investigation, Supervision, Project administration, Funding acquisition, Writing – review & editing.

## References

1. Paul BD, Snyder SH. H_2_S: A novel gasotransmitter that signals by sulfhydration. Trends Biochem Sci. 2015;40(11):687–700. doi: 10.1016/j.tibs.2015.08.007. PubMed PMID: WOS:000364728200007.

2. Luhachack L, Nudler E. Bacterial gasotransmitters: an innate defense against antibiotics. Curr Opin Microbiol. 2014;21:13–7. doi: 10.1016/j.mib.2014.06.017. PubMed PMID: WOS:000345543700004.

3. Paul BD, Snyder SH. Gasotransmitter hydrogen sulfide signaling in neuronal health and disease. Biochem Pharmacol. 2018;149:101–9. doi: 10.1016/j.bcp.2017.11.019. PubMed PMID: WOS:000429186000009.

4. Shatalin K, Shatalina E, Mironov A, Nudler E. H_2_S: a universal defense against antibiotics in bacteria. Science. 2011;334(6058):986–90. doi: 10.1126/science.1209855. PubMed PMID: WOS:000297101800058.

5. Toliver-Kinsky T, Cui WH, Törö G, Lee SJ, Shatalin K, Nudler E, et al. H_2_S, a bacterial defense mechanism against the host immune response. Infect Immun. 2019;87(1). doi: 10.1128/iai.00272-18. PubMed PMID: WOS:000453781400003.

6. Shatalin K, Nuthanakanti A, Kaushik A, Shishov D, Peselis A, Shamovsky I, et al. Inhibitors of bacterial H_2_S biogenesis targeting antibiotic resistance and tolerance. Science. 2021;372(6547):1169-+. doi: 10.1126/science.abd8377. PubMed PMID: WOS:000662088000021.

7. Teoh WP, Chen X, Laczkovich I, Alonzo F. *Staphylococcus aureus* adapts to the host nutritional landscape to overcome tissue-specific branched-chain fatty acid requirement. Proc Natl Acad Sci U S A. 2021;118(13). doi: 10.1073/pnas.2022720118. PubMed PMID: WOS:000637394200059.

8. Thurlow LR, Hanke ML, Fritz T, Angle A, Aldrich A, Williams SH, et al. *Staphylococcus aureus* biofilms prevent macrophage phagocytosis and attenuate inflammation *in vivo*. J Immunol. 2011;186(11):6585–96. doi: 10.4049/jimmunol.1002794. PubMed PMID: WOS:000290755700059.

9. Soutourina O, Poupel O, Coppée JY, Danchin A, Msadek T, Martin-Verstraete I. CymR, the master regulator of cysteine metabolism in *Staphylococcus aureus*, controls host sulphur source utilization and plays a role in biofilm formation. Mol Microbiol. 2009;73(2):194–211. doi: 10.1111/j.1365-2958.2009.06760.x. PubMed PMID: WOS:000267883900006.

10. Mironov A, Seregina T, Nagornykh M, Luhachack LG, Korolkova N, Lopes LE, et al. Mechanism of H_2_S-mediated protection against oxidative stress in *Escherichia coli*. Proc Natl Acad Sci U S A. 2017;114(23):6022–7. doi: 10.1073/pnas.1703576114. PubMed PMID: WOS:000402703800065.

11. Tikhomirova A, Rahman MM, Kidd SP, Fererro RL, Roujeinikova A. Cysteine and resistance to oxidative stress: implications for virulence and antibiotic resistance. Trends Microbiol. 2024;32(1):93–104. doi: 10.1016/j.tim.2023.06.010. PubMed PMID: WOS:001166036000001.

12. Nonoyama S, Maeno S, Gotoh Y, Sugimoto R, Tanaka K, Hayashi T, et al. Increased intracellular H_2_S levels enhance iron uptake in *Escherichia coli*. Mbio. 2024;15(10). doi: 10.1128/mbio.01991-24. PubMed PMID: WOS:001320633700001.

13. Zhang RY, Bu YF, Zhang YX, Choi SH, Wang QY, Ma Y, et al. Fur-mediated regulation of hydrogen sulfide synthesis, stress response, and virulence in *Edwardsiella piscicida*. Microbiol Res. 2024;284. doi: 10.1016/j.micres.2024.127735. PubMed PMID: WOS:001235059000001.

14. Kabil O, Banerjee R. Redox biochemistry of hydrogen sulfide. J Biol Chem. 2010;285(29):21903–7. doi: 10.1074/jbc.R110.128363. PubMed PMID: WOS:000279702200001.

15. Forte E, Borisov VB, Falabella M, Colaço HG, Tinajero-Trejo M, Poole RK, et al. The terminal oxidase cytochrome *bd* promotes sulfide-resistant bacterial respiration and growth. Sci Rep. 2016;6. doi: 10.1038/srep23788. PubMed PMID: WOS:000373168300001.

16. Forte E, Giuffrè A. How bacteria breathe in hydrogen sulfide-rich environments. Biochemist. 2016;38(5):8–11. doi: 10.1042/BIO03805008 %J The Biochemist.

17. Hammer ND, Reniere ML, Cassat JE, Zhang YF, Hirsch AO, Hood MI, et al. Two heme-dependent terminal oxidases power *Staphylococcus aureus* organ-specific colonization of the vertebrate host. Mbio. 2013;4(4). doi: 10.1128/mBio.00241-13. PubMed PMID: WOS:000326881100005.

18. Chateau A, Alpha-Bazin B, Armengaud J, Duport C. Heme a synthase deficiency affects the ability of *Bacillus cereus* to adapt to a nutrient-limited environment. Int J Mol Sci. 2022;23(3). doi: 10.3390/ijms23031033. PubMed PMID: WOS:000759368500001.

19. Svensson B, Hederstedt L. *Bacillus-subtilis*-CtaA is a heme-containing membrane-protein involved in heme-A biosynthesis. J Bacteriol. 1994;176(21):6663–71. doi: 10.1128/jb.176.21.6663-6671.1994. PubMed PMID: WOS:A1994PN64400032.

20. Xu T, Han J, Zhang J, Chen J, Wu N, Zhang W, et al. Absence of protoheme IX farnesyltransferase CtaB causes virulence attenuation but enhances pigment production and persister survival in MRSA. Front Microbiol. 2016;7. doi: 10.3389/fmicb.2016.01625. PubMed PMID: WOS:000388631700001.

21. Sun JK, Wang X, Gao Y, Li SY, Hu ZW, Huang Y, et al. H_2_S scavenger as a broad-spectrum strategy to deplete bacteria-derived H_2_S for antibacterial sensitization. Nat. Commun. 2024;15(1). doi: 10.1038/s41467-024-53764-7. PubMed PMID: WOS:001346309700016.

22. Croppi G, Zhou YY, Yang R, Bian YF, Zhao MT, Hu YT, et al. Discovery of an inhibitor for bacterial 3-mercaptopyruvate sulfurtransferase that synergistically controls bacterial survival. Cell Chem Biol. 2020;27(12):1483-+. doi: 10.1016/j.chembiol.2020.10.012. PubMed PMID: WOS:000600019000004.

23. Han TS, Yamada-Mabuchi M, Zhao G, Li L, Liu G, Ou HY, et al. Recognition and cleavage of 5-methylcytosine DNA by bacterial SRA-HNH proteins. Nucleic Acids Res. 2015;43(2):1147–59. doi: 10.1093/nar/gku1376. PubMed PMID: WOS:000350209000045.

24. Chang J, Chen B, Du ZQ, Zhao BW, Li JH, Li ZY, et al. Eugenol targeting CrtM inhibits the biosynthesis of staphyloxanthin in *Staphylococcus aureus*. Food Sci Hum Wellness. 2024;13(3):1368–77. doi: 10.26599/fshw.2022.9250115. PubMed PMID: WOS:001194808500004.

25. Li M, Rigby K, Lai YP, Nair V, Peschel A, Schittek B, et al. *Staphylococcus aureus* mutant screen reveals interaction of the human antimicrobial peptide dermcidin with membrane phospholipids. Antimicrob Agents Chemother. 2009;53(10):4200–10. doi: 10.1128/aac.00428-09. PubMed PMID: WOS:000270020600020.

26. Sun YJ, Liu MM, Niu MZ, Zhao X. Phenotypic switching of *Staphylococcus aureus* Mu50 into a large colony variant enhances heritable resistance against β-Lactam antibiotics. Front Microbiol. 2021;12. doi: 10.3389/fmicb.2021.709841. PubMed PMID: WOS:000711008300001.

27. Chen BF, Li W, Lv C, Zhao MM, Jin HW, Jin HF, et al. Fluorescent probe for highly selective and sensitive detection of hydrogen sulfide in living cells and cardiac tissues. Analyst. 2013;138(3):946–51. doi: 10.1039/c2an36113b. PubMed PMID: WOS:000312944400032.

28. Zheng JX, Shang YP, Wu Y, Wu JF, Chen JW, Wang ZW, et al. Diclazuril inhibits biofilm formation and hemolysis of *Staphylococcus aureus*. ACS Infect Dis. 2021;7(6):1690–701. doi: 10.1021/acsinfecdis.1c00030. PubMed PMID: WOS:000662225900033.

29. Guo D, Wang S, Li JH, Bai FT, Yang YP, Xu YF, et al. The antimicrobial activity of coenzyme Q0 against planktonic and biofilm forms of *Cronobacter sakazakii*. Food Microbiol. 2020;86. doi: 10.1016/j.fm.2019.103337. PubMed PMID: WOS:000495113100031.

30. Ferro TAF, Araújo JMM, Pinto BLD, dos Santos JS, Souza EB, da Silva BLR, et al. Cinnamaldehyde inhibits Staphylococcus aureus virulence factors and protects against infection in a Galleria mellonella model. Frontiers Microbiol. 2021;11. doi: 10.3389/fmicb.2020.628074. PubMed PMID: WOS:000612811300001.

31. Yang YP, Li JH, Yin Y, Guo D, Jin T, Guan N, et al. Antibiofilm activity of coenzyme Q0 against *Salmonella* Typhimurium and its effect on adhesion-invasion and survival-replication. Appl Microbiol Biotechnol. 2019;103(20):8545–57. doi: 10.1007/s00253-019-10095-8. PubMed PMID: WOS:000491439700023.

32. Nurhartadi E, Rodtong S, Thumanu K, Park SH, Aluko RE, Yongsawatdigul J. Antibacterial activity of enzymatic corn gluten meal hydrolysate and ability to inhibit *Staphylococcus aureus* in ultra-high temperature processed milk. Food Control. 2025;169. doi: 10.1016/j.foodcont.2024.110998. PubMed PMID: WOS:001355533100001.

33. Hu MJ, Zhang YY, Huang XZ, He M, Zhu JY, Zhang ZF, et al. PhoPQ regulates quinolone and cephalosporin resistance formation in *Salmonella* Enteritidis at the transcriptional level. mBio. 2023;14(3):e03395–22. doi: 10.1128/mbio.03395-22. PubMed PMID: FSTA:2024-01-Cd0405.

34. Truong-Bolduc QC, Wang Y, Ferrer-Espada R, Reedy JL, Martens AT, Goulev Y, et al. *Staphylococcus aureus* AbcA transporter enhances persister formation under β-lactam exposure. Antimicrob Agents and Chemother. 2024;68(3). doi: 10.1128/aac.01340-23. PubMed PMID: WOS:001163531600001.

35. Chang JA, Lee CY, Kim I, Kim J, Kim JH, Yun T, et al. Environmental cues in different host niches shape the survival fitness of *Staphylococcus aureus*. Nat Commun. 2025;16(1). doi: 10.1038/s41467-025-62292-x. PubMed PMID: WOS:001539302600024.

36. Peng H, Zhang YX, Palmer LD, Kehl-Fie TE, Skaar EP, Trinidad JC, et al. Hydrogen sulfide and reactive sulfur species impact proteome S-sulfhydration and global virulence regulation in *Staphylococcus aureus*. Acs Infect Dis. 2017;3(10):744–55. doi: 10.1021/acsinfecdis.7b00090. PubMed PMID: WOS:000413179000008.

37. Lan LF, Cheng A, Dunman PM, Missiakas D, He C. Golden pigment production and virulence gene expression are affected by metabolisms in *Staphylococcus aureus*. J Bacteriol. 2010;192(12):3068–77. doi: 10.1128/jb.00928-09. PubMed PMID: WOS:000278102000013.

38. Tanous C, Soutourina O, Raynal B, Hullo MF, Mervelet P, Gilles AM, et al. The CymR regulator in complex with the enzyme CysK controls cysteine metabolism in *Bacillus subtilis*. J Bio Chem. 2008;283(51):35551–60. doi: 10.1074/jbc.M805951200. PubMed PMID: WOS:000261687900027.

39. Mendes SS, Miranda V, Saraiva LM. Hydrogen sulfide and carbon monoxide tolerance in bacteria. Antioxidants. 2021;10(5). doi: 10.3390/antiox10050729. PubMed PMID: WOS:000653367700001.

40. Lensmire JM, Dodson JP, Hsueh BY, Wischer MR, Delekta PC, Shook JC, et al. The *Staphylococcus aureus* Cystine Transporters TcyABC and TcyP facilitate nutrient sulfur acquisition during infection. Infect Immun. 2020;88(3). doi: 10.1128/iai.00690-19. PubMed PMID: WOS:000514850300011.

41. Borisov VB, Forte E. Impact of hydrogen sulfide on mitochondrial and bacterial bioenergetics. Int J of Mol Sci. 2021;22(23). doi: 10.3390/ijms222312688. PubMed PMID: WOS:000742958300001.

42. Britton TA, Wu CG, Chen YW, Franklin D, Chen YM, Camacho MI, et al. The respiratory enzyme complex Rnf is vital for metabolic adaptation and virulence in *Fusobacterium nucleatum*. mBio. 2024;15(1). doi: 10.1128/mbio.01751-23. PubMed PMID: WOS:001118956600001.

43. Quirós PM, Mottis A, Auwerx J. Mitonuclear communication in homeostasis and stress. Nature Reviews Mol Cell Biol. 2016;17(4):213–26. doi: 10.1038/nrm.2016.23. PubMed PMID: WOS:000372507200007.

44. Cardamone MD, Tanasa B, Cederquist CT, Huang JW, Mahdaviani K, Li WB, et al. Mitochondrial retrograde signaling in mammals is mediated by the transcriptional cofactor GPS2 via direct mitochondria-to-nucleus translocation. Mol Cell. 2018;69(5):757-+. doi: 10.1016/j.molcel.2018.01.037. PubMed PMID: WOS:000426460900006.

45. Makhlin J, Kofman T, Borovok I, Kohler C, Engelmann S, Cohen G, et al. *Staphylococcus aureus* ArcR controls expression of the arginine deiminase operon. J Bacteriol. 2007;189(16):5976–86. doi: 10.1128/jb.00592-07. PubMed PMID: WOS:000248584800021.

46. Novák L, Zubácová Z, Karnkowska A, Kolisko M, Hroudová M, Stairs CW, et al. Arginine deiminase pathway enzymes: evolutionary history in metamonads and other eukaryotes. BMC Evol Biol. 2016;16. doi: 10.1186/s12862-016-0771-4. PubMed PMID: WOS:000386024600001.

47. Dmitriev A, Chen XR, Paluscio E, Stephens AC, Banerjee SK, Vitko NP, et al. The intersection of the *Staphylococcus aureus* Rex and SrrAB regulons: an example of metabolic evolution that maximizes resistance to immune radicals. mBio. 2021;12(6). doi: 10.1128/mBio.02188-21. PubMed PMID: WOS:000736927600001.

48. Somerville GA, Proctor RA. At the Crossroads of bacterial metabolism and virulence factor synthesis in *Staphylococci*. Microbial Mol Biol Rev. 2009;73(2):233–48. doi: 10.1128/mmbr.00005-09. PubMed PMID: WOS:000266517700002.

49. Wang R. Hydrogen Sulfide: The third gasotransmitter in biology and medicine. Antioxid Redox Signal. 2010;12(9):1061–4. doi: 10.1089/ars.2009.2938. PubMed PMID: WOS:000276421200001.

50. Eberhardt M, Dux M, Namer B, Miljkovic J, Cordasic N, Will C, et al. H_2_S and NO cooperatively regulate vascular tone by activating a neuroendocrine HNO-TRPA1-CGRP signalling pathway. Nat Commun. 2014;5. doi: 10.1038/ncomms5381. PubMed PMID: WOS:000340617600001.

51. Altaany Z, Moccia F, Munaron L, Mancardi D, Wang R. Hydrogen sulfide and endothelial dysfunction: relationship with nitric oxide. Curr Med Chem. 2014;21(32):3646–61. doi: 10.2174/0929867321666140706142930. PubMed PMID: WOS:000343166500004.

52. Lo Faro ML, Fox B, Whatmore JL, Winyard PG, Whiteman M. Hydrogen sulfide and nitric oxide interactions in inflammation. Nitric Oxide. 2014;41:38–47. doi: 10.1016/j.niox.2014.05.014. PubMed PMID: WOS:000341674800005.

53. Kolluru GK, Shen XG, Kevil CG. A tale of two gases: NO and H_2_S, foes or friends for life? Redox Biol. 2013;1(1):313–8. doi: 10.1016/j.redox.2013.05.001. PubMed PMID: WOS:000209317900043.

54. Vitko NP, Grosser MR, Khatri D, Lance TR, Richardson AR. Expanded glucose import capability affords *Staphylococcus aureus* optimized glycolytic flux during infection. mBio. 2016;7(3). doi: 10.1128/mBio.00296-16. PubMed PMID: WOS:000383440300011.

55. Beam JE, Wagner NJ, Lu KY, Fowler JG, Fowler VG, Jr., Rowe SE, et al. Inflammasome-mediated glucose limitation induces antibiotic tolerance in *Staphylococcus aureus*. Iscience. 2023;26(10). doi: 10.1016/j.isci.2023.107942. PubMed PMID: WOS:001145616900001.

